# Molecular basis of *FAAH-OUT*-associated human pain insensitivity

**DOI:** 10.1101/2022.10.20.513066

**Authors:** Hajar Mikaeili, Abdella M. Habib, Charlix Yeung, Sonia Santana-Varela, Ana P. Luiz, Kseniia Panteleeva, Sana Zuberi, Alkyoni Athanasiou-Fragkouli, Henry Houlden, John N. Wood, Andrei L. Okorokov, James J. Cox

## Abstract

Chronic pain affects millions of people worldwide. Studying pain insensitive individuals helps to identify novel analgesic strategies. Here we report how the recently discovered *FAAH-OUT* lncRNA-encoding gene, which was found from studying a pain insensitive patient with reduced anxiety and fast wound healing, regulates the adjacent key endocannabinoid system gene *FAAH*, which encodes the anandamide-degrading fatty acid amide hydrolase enzyme. We demonstrate that the disruption in *FAAH-OUT* lncRNA transcription leads to DNMT1-dependent DNA methylation within the *FAAH* promoter. In addition, *FAAH-OUT* contains a conserved regulatory element, FAAH-AMP, that acts as an enhancer for *FAAH* expression. Furthermore, using transcriptomic analyses we have uncovered a network of genes that are dysregulated from disruption of the *FAAH-FAAH-OUT* axis, thus providing a coherent mechanistic basis to understand the human phenotype observed and a platform for development of future gene and small molecule therapies.

## Introduction

Millions of people world-wide are living in chronic pain (Vos *et al*, 2017). To compound the problem, the over-prescription of opioid-based drugs to treat pain has contributed to an opioid epidemic that is causing significant morbidity and mortality, particularly in the United States (Volkow & Koroshetz, 2019). In the UK, chronic pain affects up to 50% of adults and about 12% of those have moderate-to-severe disabling pain (Mills *et al*, 2019). This has been further aggravated by the Covid-19 pandemic with up to 2 million people in the UK experiencing “long Covid” symptoms that include pain, depression and anxiety (Crook *et al*, 2021). Poorly treated chronic pain therefore makes life intolerable for extreme numbers of people and new pain-killing medications are hence urgently needed.

The endogenous cannabinoid (endocannabinoid, (eCB system or eCBS)) affects a diverse array of key physiological functions including anxiety and stress responses, pain modulation, learning and memory, wound healing and development (Lowe *et al*, 2021). It comprises the CB1 and CB2 G protein-coupled cannabinoid receptors, eCB lipid ligands (anandamide (AEA) and 2-arachidonoylglycerol (2-AG)) and their synthesising (e.g. N-acyl phosphatidylethanolamine phospholipase D (NAPE-PLD)) and metabolising (fatty acid amide hydrolase (FAAH) and monoacylglycerol lipase (MAGL)) enzymes (Cristino *et al*, 2020). The expanded eCBS includes oleoylethanolamide (OEA) and palmitoylethanolamide (PEA) lipid mediators, their receptors (e.g. TRPV1 and PPARα) and metabolic enzymes. Components of the eCBS are potential therapeutic targets for a wide range of neurological conditions including chronic pain, anxiety and depression, as well as neurodegenerative conditions such as Alzheimer’s and Parkinson’s diseases (Lowe *et al*., 2021).

A key target in the eCBS is fatty acid amide hydrolase, an important catabolic enzyme that degrades AEA, OEA, PEA and other lipids such as N-acyltaurines (NATs) (Cravatt *et al*, 1996; Deutsch & Chin, 1993; Saghatelian *et al*, 2006). FAAH is particularly enriched in the liver, brain and also expressed within trigeminal and dorsal root ganglia (DRG) (Egertova *et al*, 2003; Lau *et al*, 2014; Lever *et al*, 2009; Tsou *et al*, 1998). Within the brain, *FAAH* is found in regions that are significant for nociceptive transmission and modulation including the thalamus, periaqueductal gray (PAG) and amygdala. *Faah* is expressed in approximately a third of rat DRG neurons, with about 70% of these being TRPV1-positive (Lever *et al*., 2009). Following sciatic nerve axotomy, expression of *Faah* is also induced in large diameter DRG neurons (Lever *et al*., 2009). Over the past 20 years many FAAH inhibiting drugs have been developed, although none has yet successfully reached the clinic after human trials (Ahn *et al*, 2011; Bisogno & Maccarrone, 2013; Huggins *et al*, 2012; Tripathi, 2020). Unfortunately, a lethal toxic cerebral syndrome was precipitated by a recently trialled FAAH inhibitor (BIA 10-2474) that was later shown to be related to off-target effects (Kerbrat *et al*, 2016; van Esbroeck *et al*, 2017).

A powerful way to identify novel human-validated analgesic drug targets is to study rare individuals with intact damage-sensing neurons that present with a congenital pain insensitive phenotype (Cox *et al*, 2020). Recently we reported a new pain insensitivity disorder after studying a female patient (PFS) who, in addition to being pain insensitive, also presented with additional clinical symptoms including a happy, non-anxious disposition, fast wound healing, reduced stress and fear symptoms, mild memory deficits and significant post-operative nausea and vomiting induced by morphine (Habib *et al*, 2019). This phenotype was consistent with enhanced eCB signalling and genetic analyses showed two distinct mutations: (i) a microdeletion in a DRG and brain-expressed long non-coding RNA (lncRNA)-expressing pseudogene, *FAAH-OUT*, which is adjacent to the *FAAH* gene on human chromosome 1; and (ii) a common functional single-nucleotide polymorphism in *FAAH,* conferring reduced *FAAH* expression and activity (Chiang *et al*, 2004; Sadhasivam *et al*, 2015). These mutations result in enhanced levels of anandamide and other bioactive lipids, that are normally degraded by FAAH (Habib *et al*., 2019).

Despite *FAAH* being a heavily researched gene, the *FAAH-OUT* gene locus and how it regulates *FAAH* expression have been overlooked. Here we set up to elucidate how the ∼8kb microdeletion that is distinct from and begins ∼5 kb downstream of the 3’ end of the currently annotated footprint of the *FAAH* gene disrupts its function. Potential key mechanisms we considered included (1) the microdeleted genomic sequence contains important regulatory elements needed for normal *FAAH* expression (e.g. an enhancer); (2) the *FAAH-OUT* lncRNA transcript has an epigenetic/transcriptional role in regulating *FAAH* expression.

Here we show by gene editing in human cells that the ∼8 kb region that is deleted in PFS results in reduced expression of *FAAH*. We also demonstrate that the *FAAH-OUT* lncRNA is enriched in nuclei and its transcription positively correlates with expression of *FAAH,* bearing all the trademarks of a positive regulator. The reduction in *FAAH-OUT* transcription leads to enhanced DNA Methyltransferase 1 (DNMT1)-dependent DNA methylation of the CpG island within the *FAAH* gene promoter, resulting in transcriptional shutdown of *FAAH. FAAH-OUT* therefore appears to regulate *FAAH* expression via preventing DNMT1-dependent DNA methylation of the *FAAH* promoter, thus maintaining its transcriptional potential.

Furthermore, we show that the *FAAH-OUT* microdeletion region contains a conserved regulatory element within the first intron of *FAAH-OUT*, FAAH-AMP, that behaves as an active enhancer regulating *FAAH* expression. Editing or silencing the *FAAH-OUT* promoter region or the short evolutionarily conserved FAAH-AMP element leads to reduced *FAAH* mRNA in human cells.

Finally, to narrow in on the key functional targets downstream of the *FAAH−FAAH-OUT* axis, we used microarray analysis of patient PFS-derived fibroblasts to uncover a network of key molecular pathways and genes that become dysregulated as a result of activity disabling mutations in the *FAAH and FAAH-OUT* genes.

## Results

### Gene editing mimicking the *FAAH-OUT* microdeletion reduces *FAAH* expression

Patient PFS carries a 8,131bp heterozygous microdeletion on chromosome 1 (hg38, chr1:46,418,743-46,426,873) that begins approximately 4.9kb downstream of the end of the *FAAH* gene (**Figure 1A**) (Habib *et al*., 2019). The microdeletion contains the first two exons and putative promoter region of *FAAH-OUT (FAAHP1;* GenBank KU950306), a novel 13-exon lncRNA that is classed as a *FAAH* pseudogene and which has a similar tissue expression profile to *FAAH* (Habib *et al*., 2019).

**Figure 1.**
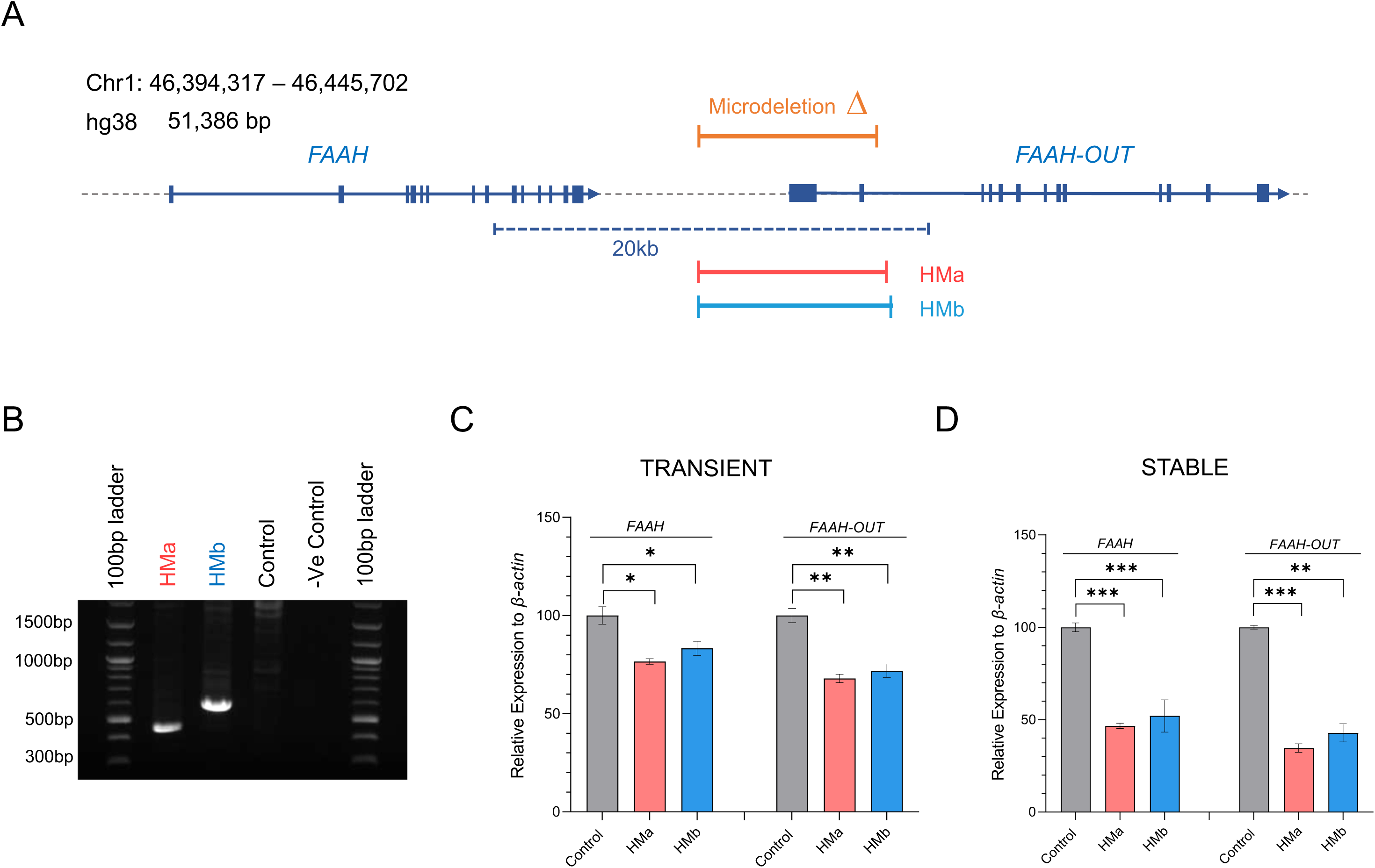
Gene editing mimicking the *FAAH-OUT* microdeletion reduces FAAH expression. A. *FAAH* and *FAAH-OUT* genomic region. Map showing human chromosome 1 (46,394,317 – 46,445,702; build hg38). *FAAH* and *FAAH-OUT* genes are shown with exons denoted by blue boxes; the direction of transcription shown by arrows. The *FAAH-OUT* gene is composed of 13 exons and the microdeletion contains the first two exons and putative promoter region. The ∼8 kb microdeletion identified in patient PFS is shown by the orange bar. Gene editing guide pairs HMa (in red) and HMb (in blue) flank the microdeleted region. **B. CRISPR/Cas9-induced microdeletion in HEK293 cells.** Gel electrophoresis of PCR products produced with primers that flank the gene editing HMa and HMb guide pairs; template genomic DNA isolated from HEK293 transiently transfected (48 hrs) with the SaCas9 plasmids. Gene editing is detected by a ∼463 bp fragment amplified from HMa edited cells and a ∼598 bp fragment from HMb edited cells. No band is observed from empty vector (control) transfected cells indicating no editing at this locus. The large size of the unedited allele is beyond the capability of the DNA polymerase. **C, D. The microdeletion in *FAAH-OUT* leads to a significant reduction in both *FAAH-OUT* and *FAAH* expression**. RT-qPCR analysis of both *FAAH-OUT* and *FAAH* mRNA levels following transient (**C**) and stable (**D**) transfections with HMa or HMb SaCas9 plasmids show significant reduction in both *FAAH-OUT* and *FAAH* expression levels. The normalized expression value of empty vector with SaCas9 but no guide RNA was set to 100, and all other gene expression data were compared to that sample. Data are expressed as mean of triplicates ± SEM and analysed by Student’s t-test, * p ≤ 0.05, ** p ≤ 0.01 and *** p ≤ 0.001.

In order to elucidate the role of the *FAAH-OUT* microdeletion on *FAAH* gene expression, we used the CRISPR/Cas9 system to edit human embryonic kidney cell lines (HEK293) to mimic the patient’s microdeletion. HEK293 cells were transiently transfected for 48 hrs with an SaCas9 plasmid bearing a guide pair (HMa or HMb, **Supplementary Table S1**) that targets sequences flanking the microdeletion, with each showing the expected genomic deletion (**Figure 1B**). Next, total RNA was isolated from the transiently transfected HEK293 cells (thus a mixture of transfected and untransfected cells) and reverse transcribed into cDNA. Quantitative real-time PCR showed a significant reduction in both *FAAH-OUT* and *FAAH* mRNAs for cells transfected with each set of guide pairs (HMa and HMb) that flank the microdeletion, highlighting that *FAAH* expression is affected by the induced downstream deletion (**Figure 1C**).

We repeated the gene editing experiments making stable HEK293 cell lines heterozygous for the *FAAH-OUT* microdeletion by transfecting SaCas9-IRES-AcGFP1 DNA plasmids carrying the HMa or HMb guide pairs. GFP-positive cells were FAC sorted to single cells to generate monoclonal lineages, and the site-specific microdeletion was confirmed by genomic DNA PCR. RT-qPCR data on *FAAH* and *FAAH-OUT* expression levels in these stable cell lines heterozygous for the *FAAH-OUT* microdeletion confirmed that *FAAH* expression is affected by the induced downstream deletion with a ∼50% reduction in *FAAH* transcript detected (**Figure 1D**).

### *FAAH-OUT* transcript is enriched in the nucleus

The *FAAH-OUT* transcript is classified as a lncRNA; it lacks a conserved protein coding sequence, is more than 200 bp in length and is post-transcriptionally capped and polyadenylated (Mattick, 2018). Studying its subcellular localization is a necessary step toward understanding the nature and mechanisms of its molecular functions.

We have shown previously that *FAAH-OUT* is expressed in a wide range of human tissues, including brain and dorsal root ganglia (Habib *et al*., 2019). Here we assessed the intracellular distribution of *FAAH* and *FAAH-OUT* transcripts using a highly sensitive fluorescence *in situ* hybridization (FISH) technology - RNAscope assay and confocal microscopy. To ensure the specific detection of *FAAH* and *FAAH-OUT* transcripts, we used probes that target different regions of each transcript.

The simultaneous visualization of *FAAH* and *FAAH-OUT* transcripts in fresh-frozen (FF) and formalin-fixed paraffin-embedded (FFPE) human tissue samples (cortex, cerebellum, prostate and DRG) provided direct evidence that *FAAH* mRNA and *FAAH-OUT* lncRNA were expressed within the same cells and predominantly localized in the cytoplasm and nucleus respectively (**Figure 2A-B, Supplementary Figures S1 and S2**). *FAAH* mRNA levels were consistently highest in *NEFH*-positive neurons in human cortex and mouse DRG (**Supplementary Figures S1B and S3).** Consistent with the FISH data, sub-cellular fractionation of HEK293 cultures followed by RT-qPCR analysis demonstrated *FAAH-OUT lncRNA* is enriched in the nucleus when compared to the *FAAH* coding mRNA, which was enriched in the cytoplasmic fraction of cells (**Supplementary Figure S1D**).

**Figure 2.**
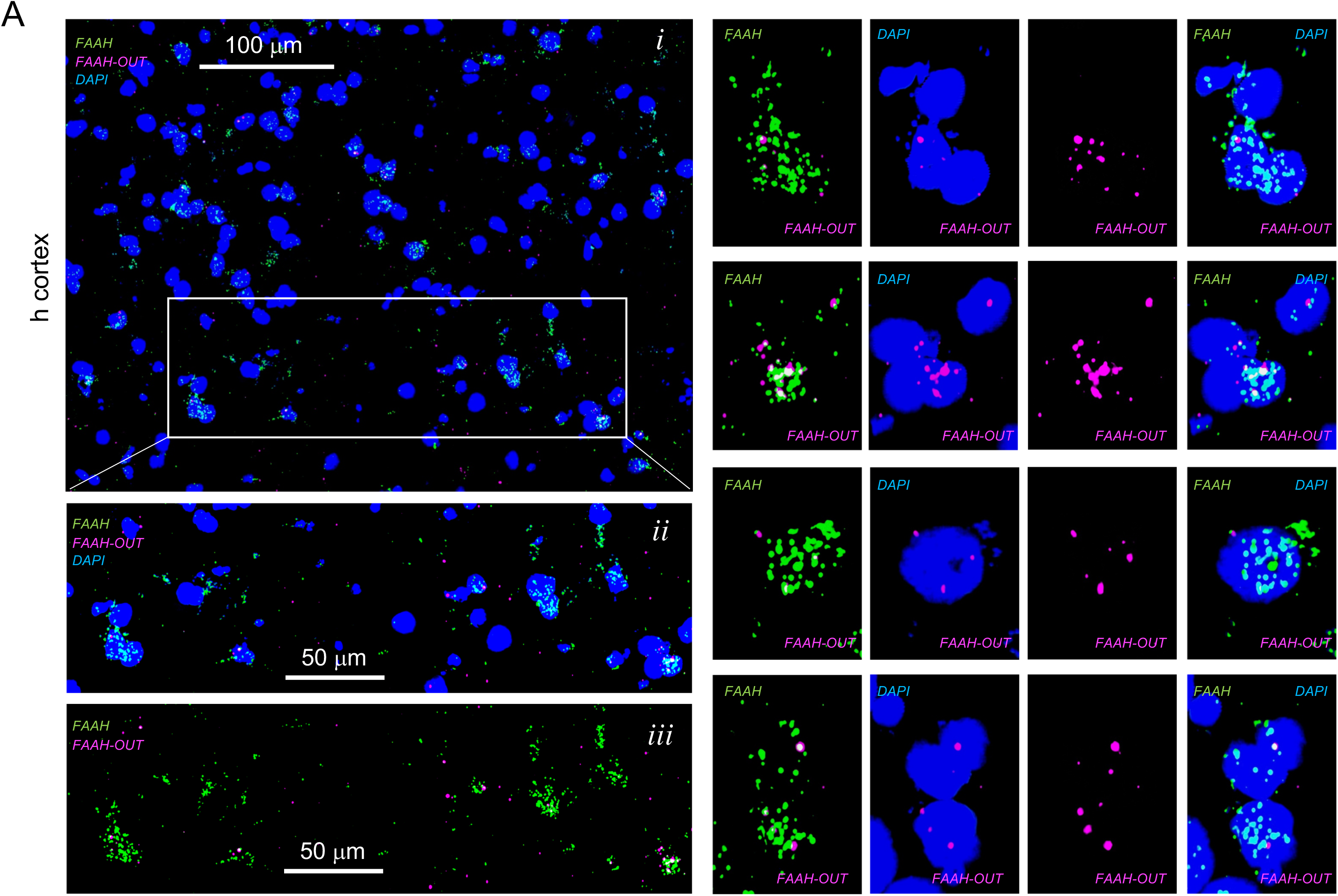
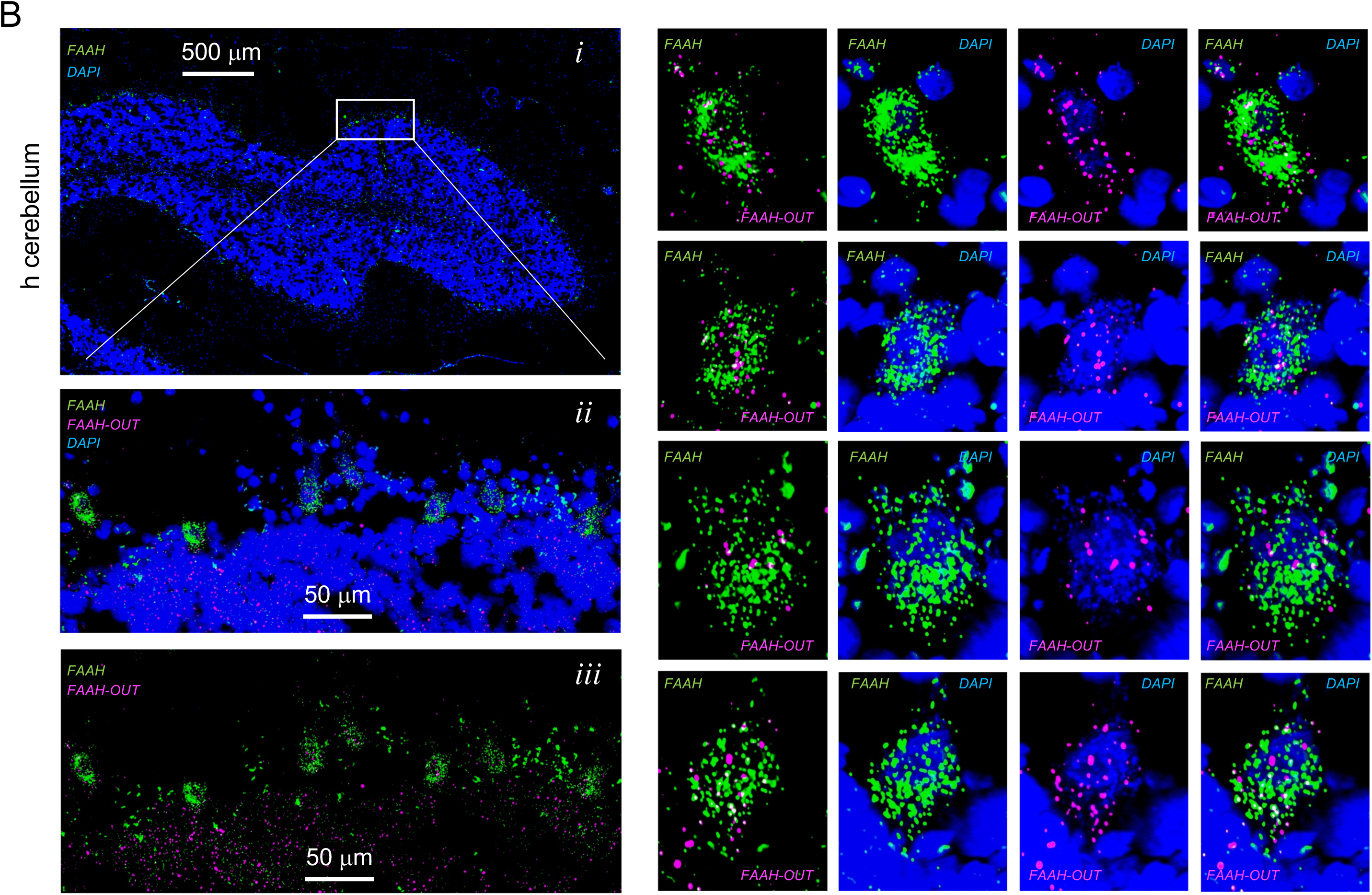
*FAAH* and *FAAH-OUT* RNA expression levels and localisation in human brain tissue cells. **A.** *FAAH* and *FAAH-OUT* RNA expression levels and localisation in human cerebral cortex cells. 7-10 μM - thick cuts of fresh frozen cerebral cortex sections were analysed by RNAscope assay (Methods). Localisation of *FAAH* mRNA (green, AF488) was compared to *FAAH-OUT* lncRNA (magenta, Opal650) localisation and DAPI staining indicating nuclei positions (blue). Scale bars are in white. A representative area (panel ***i***) and enlarged sub-area indicated by white box show colocalisation of green (*FAAH*) and magenta signal (*FAAH-OUT*) to the same cells (panels ***ii*** and ***iii***). Panels with zoomed-in individual cells expressing both *FAAH* mRNA (in green) and *FAAH-OUT lncRNA* (in magenta) are shown on the right side of the figure. **B.** *FAAH* and *FAAH-OUT* RNA expression levels and localisation in human cerebellum cells. 7-10 μM - thick cuts of fresh frozen cortex sections were analysed by RNAscope assay (Methods). Localisation of *FAAH* mRNA (green, AF488) was compared to *FAAH-OUT* lncRNA (magenta, Opal650) localisation and DAPI staining indicating nuclei positions (blue). A representative area (panel ***i***) and enlarged sub-area indicated by white box show colocalisation of green signal (*FAAH*) and magenta signal (*FAAH-OUT*) to the same large neuronal cells (Purkinje cells) located at the outer edge of cerebellar folium (panels ***ii*** and ***iii***). Panels with zoomed-in individual cells expressing both *FAAH* mRNA (in green) and *FAAH-OUT lncRNA* (in magenta) are shown on the right side of the figure and demonstrate that *FAAH* mRNA is predominantly cytoplasmic whereas *FAAH-OUT* lncRNA is enriched in the nucleus. Scale bars are in white.

### Modulation of *FAAH-OUT* transcription affects *FAAH* expression

To explore what effect *FAAH-OUT* transcription has on *FAAH* expression levels we used CRISPR/Cas9 to either (i) delete the putative *FAAH-OUT* promoter region or (ii) epigenetically silence the promoter using CRISPR interference via targeting of a nuclease-deficient form of SaCas9 (dSaCas9) fused to a Krüppel-associated box (KRAB) repressor to the *FAAH-OUT* promoter (Qi *et al*, 2013; Thakore *et al*, 2015). When localised to DNA, dSaCas9-KRAB recruits a heterochromatin-forming complex that causes histone deacetylation and methylation (H3K9 trimethylation) (Fulco *et al*, 2016; Thakore *et al*., 2015).

Guide-pair RNA sequences (**Figure 3A, Supplementary Table S1**) were selected to delete the *FAAH-OUT* promoter and cloned into an SaCas9-expressing vector. HEK293 cells were transiently transfected and the activity of each sgRNA-pair was assessed 72 hours after transfection by RT-qPCR for *FAAH* and *FAAH-OUT* mRNA expression. Both *FAAH* and *FAAH-OUT* had markedly reduced expression when the FOP2- and FOP3-guide pairs were used to induce a deletion in the *FAAH-OUT* promoter compared to cells transfected with SaCas9 only (**Figure 3B**). Similarly, epigenetic silencing of the *FAAH-OUT* promoter using the FOP1 sgRNA, which is located approximately 330 bp upstream of the transcriptional start site previously identified by 5’RACE, led to a significant reduction in *FAAH-OUT* and *FAAH* expression levels (**Figure 3C**). These results suggest that transcription of *FAAH-OUT* contributes to normal expression levels of *FAAH* and its product possibly acts as an enhancer lncRNA, similar to how *lincRNA-Cox2* functions to regulate the upstream *Ptgs2* gene (Elling *et al*, 2018).

**Figure 3.**
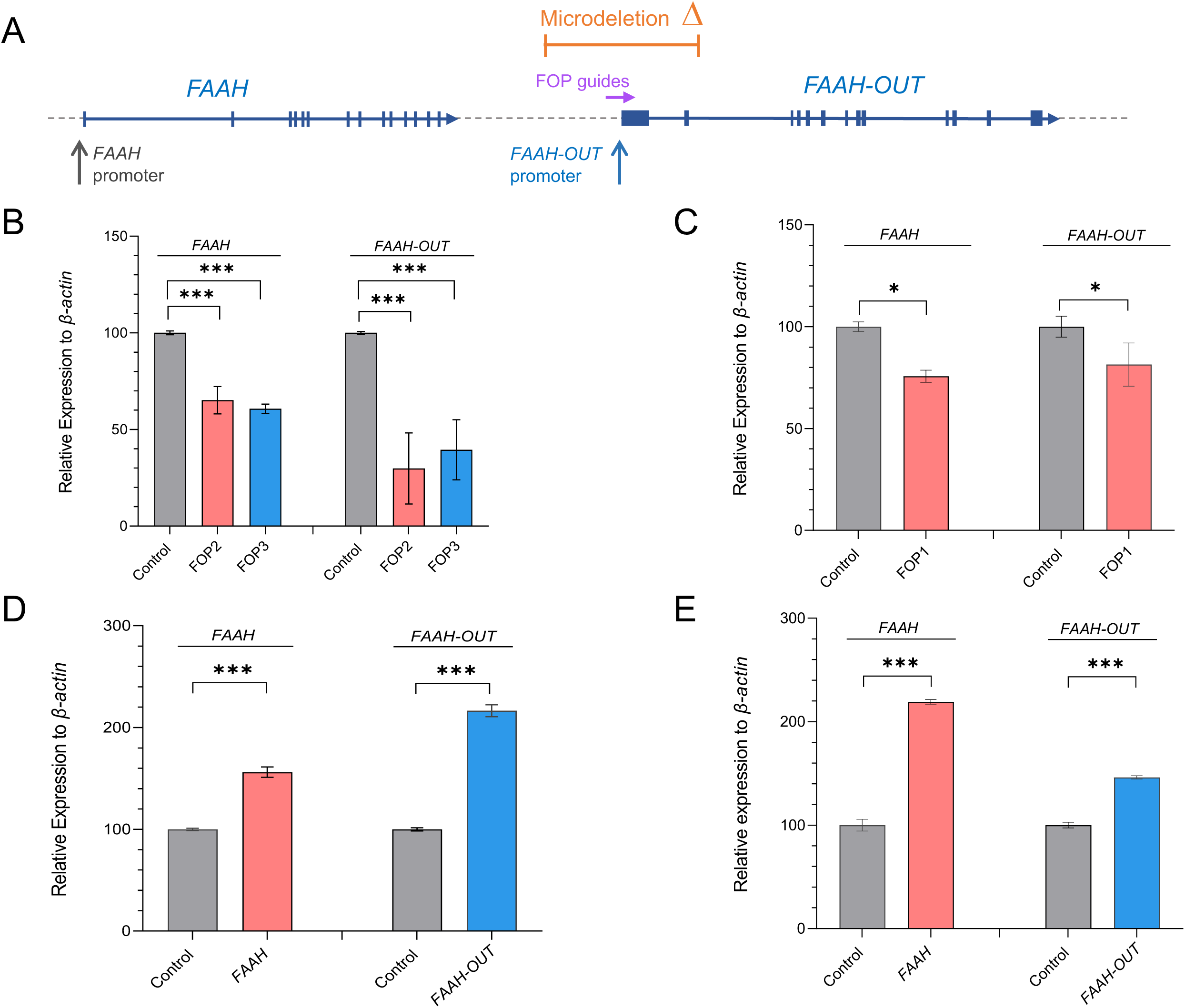
*FAAH-OUT* promoter modulates both *FAAH-OUT* and *FAAH* expression. **A.** Map showing relative positions of the ∼8 kb microdeletion identified in patient PFS (in orange), *FAAH* and *FAAH-OUT* promoters, and FOP CRISPR guides (in purple) that map to the promoter region of *FAAH-OUT.* Exons are denoted by blue boxes and the direction of transcription shown by arrows. **B. CRISPR/Cas9 -induced deletion of *FAAH-OUT* promoter leads to reduction in both *FAAH-OUT* and *FAAH* expression**. RT-qPCR analysis of both *FAAH* and *FAAH-OUT* mRNA levels showed significant reduction in both transcripts’ expression levels when HEK293 cells were transiently transfected with CRISPR/Cas9 constructs carrying either of the guide RNA pairs: FOP2 (in red) or FOP3 (in blue) designed to delete the *FAAH-OUT* promoter. **C. dSaCas9-KRAB-mediated repression of *FAAH-OUT* promoter leads to reduction in both *FAAH-OUT* and *FAAH* expression.** RT-qPCR analysis of both *FAAH* and *FAAH-OUT* mRNA levels showed significant reduction after dSaCas9-KRAB-mediated repression of the *FAAH-OUT* promoter using FOP1 guide RNA in HEK293 cells. **D. dSaCas9-VPR-mediated activation of *FAAH-OUT* promoter leads to increase in both *FAAH-OUT* and *FAAH* expression.** RT-qPCR analysis of *FAAH* and *FAAH-OUT* mRNA levels showed significant increase in both *FAAH* and *FAAH-OUT* transcript levels after targeted transcriptional activation of *FAAH-OUT* promoter using dSaCas9-VPR in HEK293 cells. dSaCas9-VPR-mediated *FAAH-OUT* activation led to transcriptional upregulation of *FAAH* gene when compared to control (empty vector). **E. dCas9-VPR-mediated activation of *FAAH* promoter leads to increase in both *FAAH* and *FAAH-OUT* expression.** RT-qPCR analysis of *FAAH* and *FAAH-OUT* mRNA levels showed significant increase in both *FAAH* and *FAAH-OUT* transcript levels after targeted transcriptional activation of *FAAH* promoter using dCas9-VPR in HEK293 cells. dCas9-VPR-mediated activation of *FAAH* also led to upregulation of *FAAH-OUT* transcript levels. In all experiments the normalized expression value of control (relevant empty vector) was set to 100, and all other gene expression data were compared to that sample. Data are expressed as mean of triplicates ± SEM and analysed by a Student’s t-test, *p ≤ 0.05, **p ≤ 0.01,***p ≤ 0.001.

To further investigate whether *FAAH-OUT* can function as an enhancer lncRNA, we employed the CRISPR activation (CRISPRa) system to recruit a strong transcriptional activator to the *FAAH-OUT* putative promoter region and activate the gene *in cis*. We successfully increased *FAAH-OUT* expression more than 2-fold in transiently transfected HEK293 cells which lead to a ∼60% increase in *FAAH* expression, as measured by RT-qPCR (**Figure 3D**). The reciprocal CRISPR activation of the *FAAH* promoter led to a more than 2-fold increase in *FAAH* mRNA levels and a ∼50% rise in *FAAH-OUT* expression (**Figure 3E**), suggesting that transcription regulation of *FAAH* and *FAAH-OUT* within this locus is interconnected.

### Highly conserved ‘FAAH-AMP’ element functions as an enhancer for *FAAH* expression

Comparative genomic analyses across species can help to identify evolutionarily conserved sequences that may have important functions (King *et al*, 2007). By analysing the PhyloP basewise conservation track for 100 vertebrates on the UCSC genome browser, a highly conserved element (denoted ‘FAAH-AMP’) was identified in the first intron of *FAAH-OUT* (**Figure 4A**). We considered that this region may contain important regulatory sequences for *FAAH-OUT* and/or *FAAH* expression.

**Figure 4.**
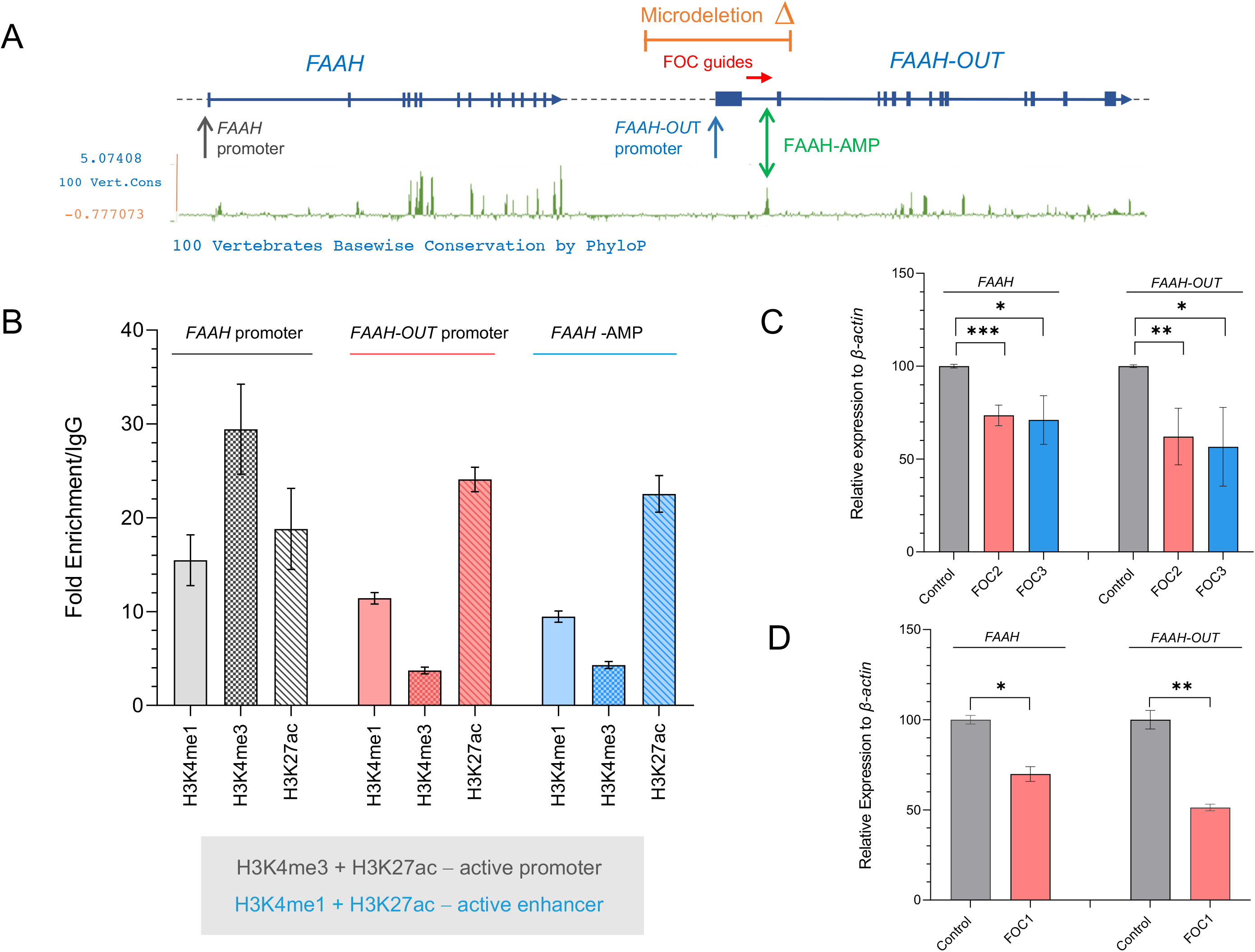
Highly conserved ‘FAAH-AMP’ element regulates both *FAAH-OUT* and *FAAH* expression. **A.** Map showing relative positions of the ∼8 kb microdeletion identified in patient PFS (in orange), *FAAH* and *FAAH-OUT* promoters, ‘FAAH-AMP’ conserved element (double green arrow) and FOC CRISPR guides (red arrow) that map to the FAAH-AMP sequence. Exons are denoted by blue boxes and the direction of transcription shown by arrows. The PhyloP base-wise conservation track for 100 vertebrates from the UCSC genome browser shows regions of high sequence conservation as peaks (in green), with the majority of these mapping to gene exons in *FAAH* and *FAAH-OUT*. **B**. **Histone modification markers at FAAH-AMP, *FAAH* and *FAAH-OUT* promoters.** ChIP-qPCR analysis at *FAAH* promoter demonstrated an enrichment of H3K4me3 and H3K27ac posttranslational modifications that typically correlate with active promoters. Both *FAAH-OUT* promoter region and FAAH-AMP element sequence were enriched in H3K4me1 and H3K27ac posttranslational modifications that typically correlate with active enhancers. **C**. **CRISPR/Cas9-induced deletion of FAAH-AMP leads to reduction in both *FAAH-OUT* and *FAAH* expression.** RT-qPCR analysis of both *FAAH* and *FAAH-OUT* mRNA levels showed significant reduction in both transcripts’ expression levels when HEK293 cells were transiently transfected with CRISPR/Cas9 constructs carrying either of guide RNA pairs: FOC2 (in red) or FOC3 (in blue) designed to delete the FAAH-AMP conserved element. **D**. **dSaCas9-KRAB-mediated repression of FAAH-AMP leads to reduction in both *FAAH-OUT* and *FAAH* expression.** RT-qPCR analysis of both *FAAH* and *FAAH-OUT* mRNA levels showed significant reduction after dSaCas9-KRAB-mediated repression of the FAAH-AMP conserved element using FOC1 guide RNA in HEK293 cells. The normalized expression value of control (relevant empty vector) was set to 100 and all other gene expression data were compared to that sample. Data are expressed as mean of triplicates ± SEM and analysed by Student’s t-test, *p ≤ 0.05, **p ≤ 0.01, ***p ≤ 0.001.

Several studies have shown that active enhancer regions are enriched in specific histone modifications including histone 3 lysine 4 (H3K4) methylation and histone 3 lysine 27 (H3K27) acetylation (Heintzman *et al*, 2009; Sethi *et al*, 2020; Shlyueva *et al*, 2014) and are highly conserved across species (Visel *et al*, 2009). We tested whether there are any enhancer marks present within the FAAH-AMP conserved region by using chromatin immunoprecipitation qPCR (ChlP-qPCR) of H3K4 and H3K27 histone marks typically associated with enhancers and promoters, including H3K4 mono-methylation (H3K4me1), H3K4 tri-methylation (H3K4me3) and H3K27 acetylation (H3K27Ac).

By comparing immuno-precipitated chromatin DNA using primers targeting either the putative FAAH-AMP enhancer region or *FAAH-OUT* putative promoter with a gene desert control region by ChIP-qPCR, we observed that both the FAAH-AMP region and the *FAAH-OUT* upstream region showed strong enrichment in H3K27ac and H3K4me1 and a low level of H3K4me3 (**Figure 4B**), a combination of post-translational modifications that is typically found at active enhancers (Buecker & Wysocka, 2012; Wang *et al*, 2015). In contrast, the *FAAH* promoter region was enriched for H3K4me3 in keeping with typical active promoter-associated histone marks (**Figure 4B**). The data therefore indicated that the FAAH-AMP conserved region indeed may function as an enhancer, potentially for *FAAH* expression.

To further test the functional importance of the FAAH-AMP conserved region as an enhancer element, we used CRISPR-Cas9 to delete the DNA containing the entire conserved FAAH-AMP region (**Figure 4A**). Targeting of SaCas9 to the FAAH-AMP region by either of two independent set of guide RNAs (FOC2 and FOC3; **Supplementary Table S1**) achieved comparable and significant reductions in *FAAH* mRNA levels confirming that the FAAH-AMP region plays a positive regulatory role for *FAAH* gene expression (**Figure 4C**).

Similarly, a reduction (∼30%) in *Faah* expression was also observed when the murine Faah-AMP region was deleted following transient transfection of mouse CAD cells with SaCas9 and the FOC4 guide-pair (**Supplementary Figure S4B**, **Supplementary Table S1**).

Next, we used a guide sequence (FOC1) to recruit dSaCas9-KRAB to FAAH-AMP to enforce inhibition of the region’s regulatory elements without cutting out the FAAH-AMP sequence. Upon transient expression of the FOC1 sgRNA with CRISPRi in HEK293 cells, we observed significant repression of *FAAH* gene expression compared to control (**Figure 4D**). Taken together these results indicate that the FAAH-AMP region contains an enhancer element that contributes to normal *FAAH* expression, and this regulatory mechanism appears to be conserved between different species. Interestingly, bioinformatic analysis of the FAAH-AMP sequence across species together with ChIP-Seq data analyses show that FAAH-AMP is a hub for transcription factor binding (**Supplementary Figure S4A**), further explaining its importance as an enhancer element.

### *FAAH-OUT* transcription modulates *FAAH* promoter methylation

Disruption of *FAAH-OUT* transcription, either by induced promoter deletion or epigenetic inhibition, leads to reduced *FAAH* expression (**Figure 3**). This can be explained by either limited access to the FAAH-AMP enhancer region and/or a potential regulatory role for the *FAAH-OUT* lncRNA. There are several reports of regulatory lncRNAs that facilitate the status of target promoter and/or enhancer regions, such as recruiting chromatin remodellers, transcription factors and DNA modifiers such as DNA methylases or DNA demethylases (Chalei *et al*, 2014; Di Ruscio *et al*, 2013; Miao *et al*, 2019; Zhao *et al*, 2016). Considering that the *FAAH* promoter has a strong and conserved CpG island, modulating DNA methylation status in order to regulate *FAAH* expression is a possibility. Furthermore, the *FAAH* gene region sequence has been reported to have DNMT1-dependent DNA methylation (Di Ruscio *et al*., 2013).

CpG-rich promoters are typically unmethylated, marked with histone modifications such as H3K4me3, and are highly active. If the *FAAH-OUT* lncRNA normally regulates levels of DNA methylation at the *FAAH* promoter, then loss of one *FAAH-OUT* allele (like in patient PFS) could be sufficient to shift the balance of DNA methylation and chromatin modification towards *FAAH* promoter inactivation.

To test whether the reduction in *FAAH-OUT* expression affects the local epigenomic profile at the *FAAH* promoter region, we used DNA methylation and ChIP-qPCR assays to screen for levels of methylated DNA at the *FAAH* promoter in a heterozygous HEK293 *FAAH-OUT*^+/-^ cell line. As shown in **Figure 5A**, in normal wild type (WT) cells, methylation levels at the *FAAH* promoter are balanced between 40% methylated and 60% unmethylated DNA, with methylation rising sharply by ∼60% in heterozygous (HTZ) cells with reduced levels of *FAAH-OUT* expression, reversing the methylated vs unmethylated ratio. This suggests that lower levels of the *FAAH-OUT* lncRNA due to one allele loss leads to local epigenetic changes that drive *FAAH* expression down. Furthermore, the epigenetic inactivation of the *FAAH* promoter is enhanced by a rise in H3K9me3 modification (**Figure 5B**).

**Figure 5.**
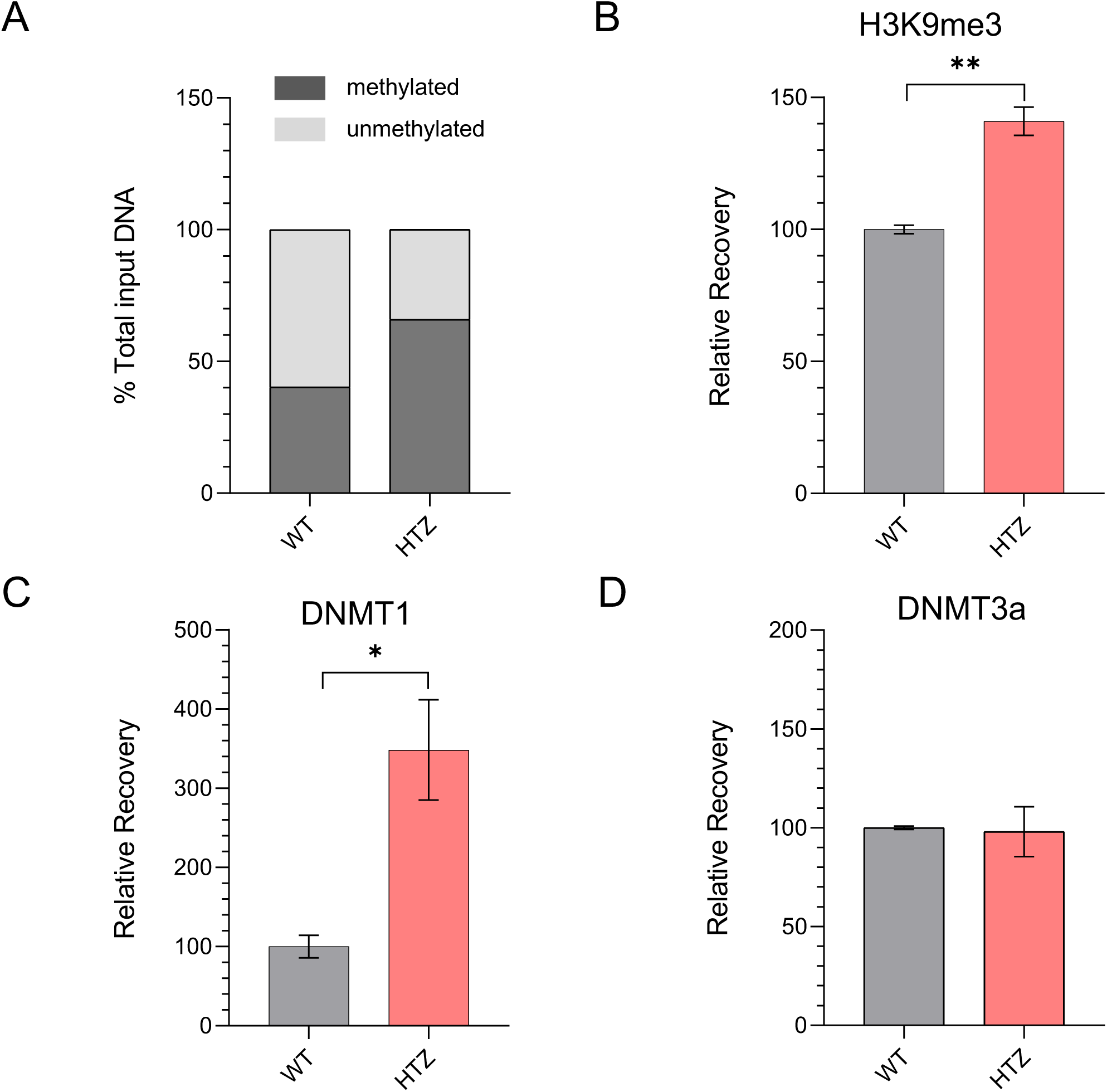
*FAAH* expression is regulated by DNA methylation in a DNMT1- and *FAAH-OUT*-dependent manner. **A**. *FAAH* promoter methylation is increased when *FAAH-OUT* levels are reduced. MethylScreen analysis showed a significant increase in *FAAH* promoter DNA methylation levels in HEK293-HTZ stable line cells harbouring heterozygous (HTZ) *FAAH-OUT* microdeletion (∼40% methylation in WT cell line to ∼65% methylation in HTZ line). **B**. ***FAAH* promoter is less active when *FAAH-OUT* transcription is reduced.** ChIP-qPCR analysis at *FAAH* promoter showed that the increased DNA methylation at the *FAAH* promoter in HEK293-HTZ cells coincides with an increase in H3K9me3 posttranslational modification that typically correlates with heterochromatin, both indicating a less active *FAAH* promoter state. **C** and **D**. **DNA at *FAAH* promoter is methylated by DNMT1.** ChIP-qPCR analysis at *FAAH* promoter in HEK293-HTZ cells showed a significant (∼3-fold) increase in chromatin-associated DNMT1 (**C**) when compared to the wild type (WT) control, whereas the levels of chromatin-bound DNMT3a did not change (**D**). Mean values from the WT control cells were assigned as 100%. Data are expressed as mean of triplicates ± SEM and analysed by Student’s t-test, *p ≤ 0.05, **p ≤ 0.01 (**B-D**).

We next explored whether this enhanced DNA methylation at the *FAAH* promoter in heterozygous HEK293 *FAAH-OUT*^+/-^ cells is provided by one of the known DNA methylases. ChIP-qPCR analysis showed that DNMT1 localisation at the *FAAH* promoter was enriched 3-fold, whereas levels of DNMT3a did not change (**Figure 5C and D**). This indicates that loss of the *FAAH-OUT* allele and/or reduction in *FAAH-OUT* lncRNA levels lead to increased recruitment of DNMT1 to the *FAAH* promoter and a rise in DNA methylation, in keeping with previously reported data for DNMT1-dependent genome-scale methylation profiling (Di Ruscio *et al*., 2013).

### Transcriptomic analyses of PFS patient-derived fibroblasts

Primary fibroblast cell lines derived from patient PFS and 4 unrelated female healthy controls were cultured and total RNA isolated. *FAAH* expression, as shown by RT-qPCR, was significantly downregulated in the patient-derived fibroblast cell line compared to controls (**Figure 6D**). To explore whether additional genes were also dysregulated and to identify potential downstream candidate genes and pathways that could help explain the PFS phenotype, we carried out a whole transcriptome microarray. This showed striking gene dysregulation (**Figure 6, Supplementary Figure S5, Supplementary Table S5**) with 797 genes upregulated and 348 genes downregulated (>2 fold change; p<0.05) between the PFS line and 4 controls. Ingenuity Pathway analyses highlighted groups of gene products which take part in WNT-induced signalling, wound healing, BDNF-signalling and G-protein signalling (**Figure 6A**). A number of genes connected to WNT-regulated pathways were dysregulated included the downregulated stimulators of canonical WNT-dependent pathway *SFRP2* and *SERPINF1,* the upregulated repressor *DKK1*, and the upregulated *WNT5B and WNT16* transcription factors (**Figure 6B, Supplementary Figure S6**) (Cadigan & Peifer, 2009; Liang *et al*, 2019; Nusse & Varmus, 2012; Suthon *et al*, 2021; van Amerongen & Nusse, 2009; Willert & Nusse, 2012).

**Figure 6.**
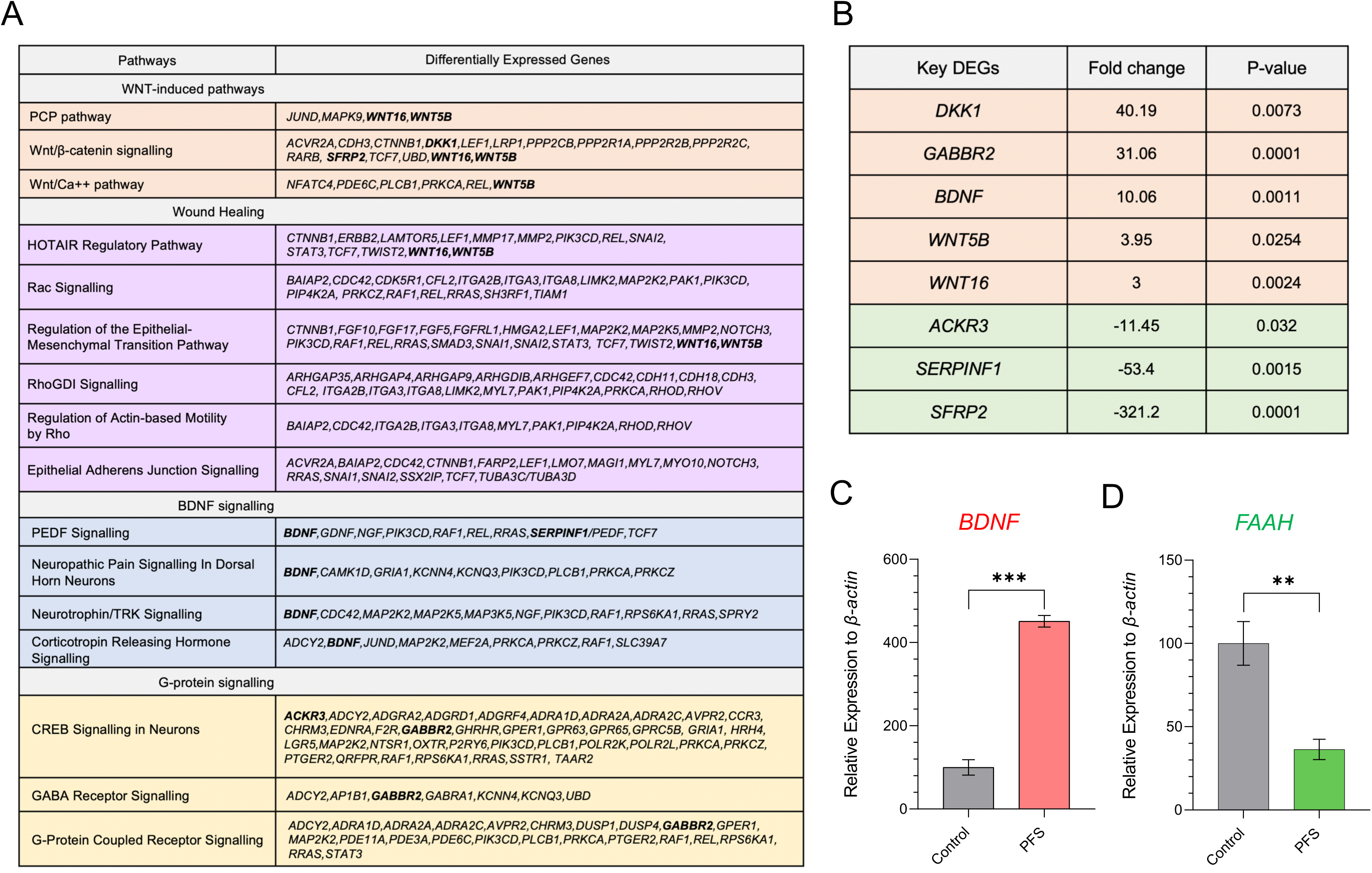
Differential gene expression in cells with *FAAH-OUT* microdeletion. Patient PFS derived fibroblast cell line and 4 gender matched control cell lines were analysed using microarrays. Selected pathways of interest that contain significantly differentially expressed genes (DEGs) are shown in (**A**), key differentially expressed genes in (**B**) and the list of DEGs can be seen in **Supplementary Table S5**. **C. *BDNF* mRNA levels rise in patient fibroblasts.** RT-qPCR analysis of *BDNF* mRNA levels in patient PFS derived fibroblasts showed a significant rise in *BDNF* mRNA levels when compared to the four gender matched controls. **D. *FAAH* mRNA levels are reduced in patient fibroblasts**. RT-qPCR analysis of *FAAH* mRNA levels in patient PFS derived fibroblasts showed significant reduction in *FAAH* mRNA levels when compared to four gender matched controls. For C and D data were normalized to the beta-actin gene as an endogenous control. The normalized expression value of control subjects was set to 100. Data are expressed as mean of triplicates ± SEM and analysed by Student’s t-test. **p ≤ 0.01, *** p ≤ 0.001

One gene of interest that was significantly upregulated in the PFS cell line was brain-derived neurotrophic factor (*BDNF*), with the microarray assay result validated by RT-qPCR (**Figure 6C**). Interestingly, previous work in rats has shown that pharmacological inhibition of the FAAH enzyme elevated BDNF levels (Carnevali *et al*, 2020; Shang *et al*, 2022; Vinod *et al*, 2012). We replicated this data in wild-type mice by showing that systemic injection of FAAH inhibitor URB597 showed a ∼25% increase in hippocampal BDNF levels, as determined by ELISA (**Supplementary Figure S7A**). The connection between loss of FAAH activity and increased BDNF levels is particularly interesting given the patient’s reported elevated mood and the known anti-depressive actions of BDNF and TrkB signalling (Casarotto *et al*, 2021; Habib *et al*., 2019).

Another gene of interest that is significantly downregulated in the patient PFS cell line encodes the atypical chemokine receptor *ACKR3*, with the microarray result validated by RT-qPCR (**Supplementary Figure S6E**). ACKR3 is a broad-spectrum opioid scavenger receptor, downregulation of which could contribute to the painless phenotype (Ehrlich *et al*, 2021; Meyrath *et al*, 2020; Szpakowska *et al*, 2021). We confirmed the connection between *FAAH* downregulation and *ACKR3* transcript levels by using silencing RNA against *FAAH* in a HEK293 cell line, which led to a ∼40% decrease in *ACKR3* levels (**Supplementary Figure S7B**).

Patient PFS has previously observed that wounds heal quickly and work carried out in mice has shown that genetic or pharmacological inhibition of FAAH activity accelerates skin wound healing (Sasso *et al*, 2016). We analysed cell migration of the PFS fibroblasts compared to control fibroblasts using a scratch assay and time-lapse microscopy which showed that gap closure was significantly faster in the PFS fibroblasts (**Supplementary Figure S8**). This is consistent with previous work where human keratinocytes also showed a marked increase in migration in a scratch assay in the presence of a FAAH inhibitor and supports FAAH as a potential therapeutic target for wound healing (Sasso *et al*., 2016).

## Discussion

In this study we provide the first mechanistic insights into how the microdeletion identified in patient PFS negatively affects *FAAH* expression and leads to pain insensitivity, accelerated wound healing and the lack of depression and anxiety symptoms observed in the patient. The ∼8kb microdeletion contains the upstream promoter region and first two exons of *FAAH-OUT* and also an evolutionary conserved ‘FAAH-AMP’ element in the first intron that contains several potential transcription factor binding sites. We show that FAAH-AMP has the chromatin marks of an active enhancer and positively regulates *FAAH* expression. CRISPR interference at the FAAH-AMP enhancer element or the *FAAH-OUT* promoter results in reduced levels of both *FAAH-OUT* and *FAAH* mRNA in human cells, indicating that *FAAH* expression is regulated by transcriptional activity at the *FAAH-OUT* locus. The FAAH-AMP enhancer element is also conserved in mice, with CRISPR editing similarly leading to a reduction in *Faah* expression. Furthermore, we show that *FAAH* and *FAAH-OUT* are co-expressed within the same cells, with the *FAAH-OUT* lncRNA being enriched in nuclei as shown by RNAscope experiments in different human tissues and cell fraction analyses.

The data suggest two mechanisms of *FAAH-OUT*-dependent *in cis* regulation of *FAAH* in which transcription of the *FAAH-OUT* gene leads to (1) expression of the *FAAH-OUT* lncRNA that may play a role as a positive regulator of *FAAH*, and (2) opening up of chromatin in the FAAH-AMP enhancer region which improves accessibility to the region for proteins that in turn modulate efficiency of *FAAH* transcription and potentially allow local looping between *FAAH* and *FAAH-OUT* genes for co-ordinated transcription (**Figure 7A** and **B**). The reduction in *FAAH-OUT* transcription leads to enhanced DNMT1-dependent DNA methylation of the CpG island within the *FAAH* gene promoter, and subsequent chromatin remodeling as witnessed by increased H3K9 trimethylation, resulting in transcriptional shutdown of *FAAH.* The *FAAH-OUT* lncRNA may therefore regulate *FAAH* expression via preventing DNMT1-dependent DNA methylation of the *FAAH* promoter, thus maintaining its transcriptional potential. DNMT1 methylation regulation of the *FAAH* promoter has previously been reported, as have examples of other lncRNAs that regulate DNA methylation at the promoter regions of other genes (Chalei *et al*., 2014; Di Ruscio *et al*., 2013; Zhao *et al*., 2016).

**Figure 7.**
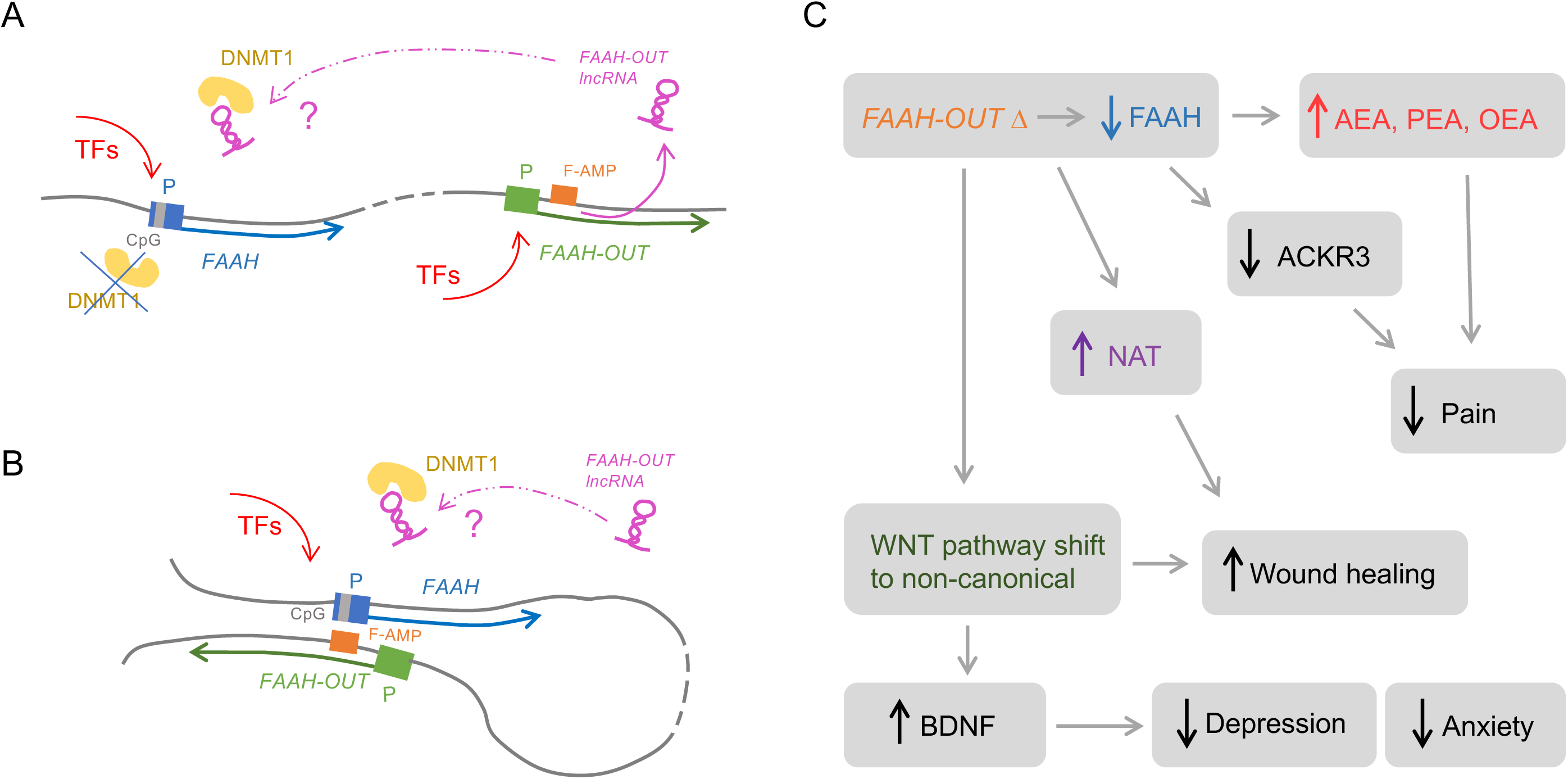
Schematic representation of *FAAH-OUT* - dependent regulation of *FAAH* expression and subsequent phenotypical changes in the patient. **A.** *FAAH-OUT* transcription regulates *FAAH* expression via preventing DNMT1-dependent DNA methylation at *FAAH* promoter. Activation of *FAAH-OUT* promoter (in green) leads to transcription of *FAAH-OUT* lncRNA (in purple) and opening of chromatin at FAAH-AMP (F-AMP, in orange)*. FAAH-OUT* lncRNA potentially prevents DNMT1 (in yellow) recruitment to methylate *FAAH* promoter (in blue) at CpG island (in grey) allowing *FAAH* gene to be transcribed at higher levels. **B.** Additional level of regulation via FAAH-AMP enhancer (F-AMP, in orange) could be provided via potential looping between the enhancer and *FAAH* promoter in blue to facilitate transcription factors (TFs) binding. **C.** Schematic network of key dysregulated genes and pathways that result from the disruption of the *FAAH−FAAH-OUT* axis, providing molecular basis for the phenotypes observed in the patient. Microdeletion in *FAAH-OUT* leads to reduction in *FAAH* expression and subsequent fall in overall FAAH activity thus leading to rise in endocannabinoid levels (AEA, PEA and OEA) which (especially anandamide, AEA) facilitate pain insensitivity. The mutation also leads to a drop in ACKR3 levels, lack of which as a broad-spectrum scavenger for opioid peptides adds another potential level to the patient’s analgesia. In addition, the decrease in FAAH activity leads to a rise in N-acyl taurine (NAT) and changes in WNT pathways (shift from canonical to non-canonical) both of which likely contribute to accelerated wound healing. The WNT pathway shift also leads to a dramatic rise in BDNF levels, thus protecting the patient from depression and anxiety.

Further work will help to understand exactly how *FAAH-OUT* may be functioning as an enhancer RNA and whether the *FAAH-OUT* lncRNA forms complexes directly at the *FAAH* promoter and/or FAAH-AMP region, similar to other known lncRNA transcriptional regulators (Chalei *et al*., 2014; Di Ruscio *et al*., 2013; Miao *et al*., 2019; Zhao *et al*., 2016). In addition to possibly protecting the *FAAH* promoter from DNMT1-dependent DNA methylation, the *FAAH-OUT* lncRNA may play a role in keeping the *FAAH* promoter active by maintaining an R-loop at that region. R-loops (three-stranded RNA/DNA structures) form when a nascent transcript or a lncRNA invades and makes a complex with a DNA duplex and are widespread at the GC-rich regions of promoters, protecting CpG islands from DNA methylation and preventing silencing (Chédin, 2016; Tan-Wong *et al*, 2019). Both *FAAH* and *FAAH-OUT* promoters are GC-rich with more than one hundred CpG pairs within the *FAAH* promoter sequence and about half of those are clustered in large CpG islands. R-loop formation could be achieved by locking RNA onto the DNA strand via 4xG repeats using a velcro-type interaction between them and quadruple Cs on the other strand (Ginno *et al*, 2012). The *FAAH* promoter sequence has about ten 4xGs whereas the *FAAH-OUT* lncRNA has twenty 4xGs.

It is also of interest that chromatin marks at the *FAAH-OUT* promoter region were similar to those at the FAAH-AMP region and resembled those of an active enhancer (enriched in H3K27Ac and H3K4me1) rather than a promoter (enriched in H3K4me3 and H3K27Ac) (**Figure 4B**). This suggests that either the *FAAH-OUT* promoter has a dual role serving as a promoter for *FAAH-OUT* and an enhancer for *FAAH* or it has been evolutionary repurposed from promoter to enhancer, thus modulating *FAAH* expression together with FAAH-AMP (Carelli *et al*, 2018; Mikhaylichenko *et al*, 2018). This would be consistent with our data where transcriptional activation of *FAAH-OUT* via the CRISPRa system leads to an upregulation of *FAAH* and vice versa, where activation of *FAAH* transcription leads to an increase in *FAAH-OUT* RNA levels (**Figure 3D** and **E**).

We further aimed to understand how decreased levels of FAAH and higher levels of anandamide (and other substrates) translate into the patient’s pain insensitivity syndrome, which is characterised by the absence of post-operative pain, painless burns and bone fractures, a happy, non-anxious disposition, fast wound healing, fear and memory deficits, and significant post-operative nausea and vomiting induced by morphine (Habib *et al*., 2019). In the original case report, we also noted that her dental surgeon observed, most unusually, that her saliva dissolves the fixative for a temporary denture after just 90 mins (Habib *et al*., 2019). Interestingly, there is a recent report of lipidomic profile differences in the submandibular gland of FAAH knockout versus wild-type mice (Andreis *et al*, 2022). It will be interesting to assay the salivary lipidomic profiles of patient PFS in potential future studies.

To narrow in on the key functional targets downstream of the *FAAH - FAAH-OUT* axis we used microarray analysis of patient-derived fibroblasts to uncover a network of key molecular pathways and genes that become dysregulated as a result of disrupting *FAAH-OUT*. There were 797 genes upregulated and 348 genes downregulated (>2 fold change; p<0.05) between the PFS line and 4 gender-matched controls. Pathway analyses showed major changes in expression level of genes which take part in WNT-induced signalling, wound healing, BDNF-signalling and G-protein signalling (**Figure 6A**). Thus, several genes connected to WNT-regulated pathways included upregulation of the *DKK1* repressor and *WNT5B and WNT16* transcription factors and downregulation of canonical WNT-dependent pathway stimulators such as *SFRP2.* This combined indicates that the reduced FAAH levels and activity lead to WNT pathway(s) shift from a canonical to non-canonical type. Importantly, WNT pathways have been previously linked to wound healing and both upregulated *Wnt5b* and *Wnt16* have also been linked to bone regeneration (Cadigan & Peifer, 2009; Gori *et al*, 2015; Kobayashi *et al*, 2015; McGowan *et al*, 2021; Suthon *et al*., 2021). In addition to gene expression changes highlighted in **Figure 6**, FAAH is known to degrade N-acyl taurines (NATs) which are lipids implicated in regulation of skin wound healing, thus further helping to explain the accelerated healing phenotype observed for the PFS patient (Sasso *et al*., 2016).

Interestingly, WNT-dependent signalling has been previously reported to be connected to levels of brain-derived neurotrophic factor (BDNF), which modulates mood and is directly linked to anxiety and depression through TrkB receptor signalling (Casarotto *et al*., 2021; Wang *et al*, 2022; Yi *et al*, 2012). Furthermore, pharmacological inhibition of FAAH activity has been reported to lead to an increase in BDNF levels in rats (Carnevali *et al*., 2020; Shang *et al*., 2022; Vinod *et al*., 2012). Our gene expression analyses in patient-derived fibroblasts show a significant upregulation in *BDNF* expression, although whether this is replicated in other patient tissues remains to be tested. Nevertheless, we have shown that in mice, pharmacological inhibition of the FAAH enzyme also upregulates hippocampal BDNF, providing further evidence for the FAAH-BDNF link.

Another gene that is significantly upregulated is *GABBR2* which encodes receptor subunit GABAB_2_ which forms an active heterodimeric complex with GABAB_1_ in the GABAB receptor (Cryan & Kaupmann, 2005). GABAB receptors are abundant in the brain, where they are localized in many neuronal cell types including interneurons and some glial cells (Cryan & Kaupmann, 2005). GABBR2 inhibits neuronal activity via G-protein coupled secondary messenger systems and its low levels were implicated in reduced analgesic effects of oxycodone (Thibault *et al*, 2014). Furthermore, GABAB receptor knockout mice data indicate a role for GABAB receptors in nociception and anxiety, with GABAB knockout mice showing increased anxiety (Cryan *et al*, 2004; Mombereau *et al*, 2004). Interestingly, low levels of GABBR2 expression were shown to be a valid biomarker for patients with chronic migraine (Plummer *et al*, 2011). Thus high levels of GABBR2 expression could be consistent with the pain- and anxiety-free phenotype of patient PFS (Habib *et al*., 2019).

Among significantly downregulated genes, one of particular interest is *ACKR3* (**Figure 6, Supplementary Figure S6**). *ACKR3* encodes the atypical chemokine receptor ACKR3/CXCR7 that is widely expressed in brain. ACKR3 has recently been reported as a broad-spectrum scavenger for opioid peptides and has also been identified as a natural target of conolidine, a natural analgesic alkaloid (Meyrath *et al*., 2020; Szpakowska *et al*., 2021). These properties potentially make ACKR3 an important and physiologically relevant contributor to the PFS patient painless phenotype. A reduction in *ACKR3* expression levels resulting from downregulation of the *FAAH-FAAH-OUT* axis could lead (via downregulation of *ACKR3*) to higher availability of endogenous opioid peptides for the classical opioid receptors (**Figure 7C**).

In summary, our data show that microdeletion in *FAAH-OUT* disrupts transcription of the *FAAH-OUT* lncRNA and eliminates the enhancer sequence element FAAH-AMP, thus leading to deregulation of the *FAAH-FAAH-OUT* axis. We demonstrate that reduction in *FAAH-OUT* transcription leads to DNMT1-dependent DNA methylation of the CpG island within the *FAAH* gene promoter, resulting in transcriptional shutdown of *FAAH* and reduction of FAAH activity. Moreover, through microarray analysis of patient PFS-derived fibroblasts we have uncovered a network of key molecular pathways and genes that become dysregulated as a result of disrupting *FAAH-OUT* such as a shift in WNT-dependent pathways towards non-canonical, a dramatic increase in *BDNF* and a decrease in *ACKR3* expression levels.

Whilst further experiments would be needed to elucidate the precise mechanism(s) by which the *FAAH-OUT* lncRNA regulates *FAAH*, our data provide a significant advance in understanding inter-pathway crosstalk resulting from lower FAAH activity and higher anandamide levels connecting for the first time major players from the endocannabinoid system with those of G-protein and opioid signaling. The data thus provide a coherent explanation for the pain insensitivity, lack of anxiety, faster wound-healing and other syndromic symptoms observed in the patient and form a platform for development of future gene and small molecule therapies. Given the current failure of small molecule inhibitors of FAAH as human analgesics, our findings validate *FAAH-OUT* regulation of *FAAH* as a new route to develop pain treatments.

## Materials and Methods

### CRISPR/Cas9 plasmids

Plasmids 61591 (Ran *et al*, 2015) and 106219 (Thakore *et al*, 2018) (Addgene) were modified for the gene editing (SaCas9) and transcriptional repression (dSaCas9-KRAB) CRISPR experiments respectively. For gene editing plasmid 61591, the CMV promoter was replaced with a shorter promoter sequence derived from the housekeeping *Eef1a1* gene and the bGH polyadenylation sequence was replaced with a shorter synthetic polyadenylation sequence. gBlocks gene fragments (Integrated DNA Technologies) were designed to contain a U6 promoter, guide sequence and modified guide scaffold (Tabebordbar *et al*, 2016) with the design enabling two guide cassettes to be inserted into one SaCas9 plasmid by In-Fusion cloning (Takara). Guide sequences were designed using the online CRISPOR design tool (Haeussler *et al*, 2016). The sgRNAs chosen were based on a high specificity rank and a low potential off-target score. The ‘empty vector’ control contained *Eef1a1*-promoter driven SaCas9 but no guide sequences.

SaCas9-IRES-AcGFP1 versions of the plasmids were generated by inserting an IRES-AcGFP1 sequence (Integrated DNA Technologies) after the SaCas9 sequence but before the polyadenylation sequence (at the AflII restriction site) by In-Fusion cloning.

For the transcriptional repression CRISPRi experiments the AgeI-EcoRI fragment from plasmid 106219 containing the dSaCas9-KRAB sequence was used to replace the SaCas9 sequence from plasmid 61591, to give a CMV driven dSaCas9-KRAB. Next, a gBlocks gene fragment (Integrated DNA Technologies) was designed to contain a synthetic poly(A) sequence, U6 promoter, guide sequence and modified guide scaffold (Tabebordbar *et al*., 2016). This sequence was cloned into the EcoRI-NotI sites of the modified plasmid 61591 using In-Fusion cloning (Takara). The ‘empty vector’ control contained CMV-promoter driven dSaCas9-KRAB but no guide sequences. All transformations were performed in Stbl3 *E. coli* strain (Thermo Fisher).

Plasmid 68495 (Kiani *et al*, 2015) (Addgene) was modified for the transcriptional activation (CRISPRa) experiments. The FseI-EcoRI fragment from plasmid 68495 containing the VP64-p65-Rta (VPR) sequence was used to replace the KRAB sequence from the CMV driven dSaCas9-KRAB plasmid to give a transcriptional activator plasmid. Next, a gBlocks gene fragment (Integrated DNA Technologies) was designed to contain a synthetic poly(A) sequence, U6 promoter, guide sequence and modified guide scaffold (Tabebordbar *et al*., 2016). This sequence was cloned into the EcoRI-NotI sites of the CRISPRa plasmid using In-Fusion cloning (Takara). All guide sequences and their genomic locations are listed in **Supplementary Table S1** and PCR primers used to verify editing are listed in **Supplementary Table S2**.

### Transient transfection of CRISPR/Cas9 plasmids into HEK293 and CAD cells

Human embryonic kidney 293 cells (HEK293, ECACC) were cultured in Dulbecco’s Modified Eagle’s Medium (Thermo Fisher) with 10% fetal bovine serum (Hyclone). Mouse catecholaminergic neuronal tumour cells (CAD, ECACC) were cultured in DMEM:HAMS F12 (1:1) with 2% glutamine and 8% fetal bovine serum. Lipofectamine 3000 (Invitrogen) was employed as a DNA carrier for transfection according to the manufacturer’s procedures. Briefly, Lipofectamine 3000 was diluted into Opti-MEM I Reduced Serum Medium (Life Technologies, Inc.). Plasmid DNA was first diluted into Opti-MEM and 10µl of P3000 reagent was added to the mixture. The DNA-liposome complex was prepared by adding diluted DNA into diluted Lipofectamine (ratio 1:1) and incubating the mixture at room temperature for 30 min. DNA-liposome mixture was added to 70% confluent HEK293/CAD cells. After 24 hours of incubation at 37 °C, media was removed and the transfection steps were repeated. The cells were incubated at 37°C in a 5% CO_2_ incubator with 92-95% humidity for another 24 hours.

To extract total RNA from cultured cells, medium was first aspirated off and cells were rinsed with ice cold PBS. 1ml of TRIzol (Invitrogen) was added directly to the cells and was incubated for 5 minutes at room temperature. Cell lysate was passed through a pipette up and down several times. RNA was extracted using PureLink™ RNA Micro Scale Kit (Invitrogen) according to the manufacturer’s procedures. Genomic DNA was isolated using the DNeasy Blood and Tissue kit (Qiagen) and used as template to confirm gene editing (**Supplementary Table S2**).

### Generation of stable cell lines

HEK293 cells were transfected using Lipofectamine 3000 with SaCas9-IRES-AcGFP1 plasmids containing guide pairs HMa or HMb (**Supplementary Table S1**) or a no guide control. For FACS, cells were washed with PBS and detached using trypsin. The cell pellets were washed twice with PBS and resuspended in ice cold PBS, 5mM EDTA, 25mM HEPES buffer and 1% FBS. The top 3% GFP-positive cells were sorted into 96-well plates (1 cell per well) and cultured for 3 weeks. Genomic DNA was isolated using the DNeasy Blood and Tissue kit (Qiagen) and screened for the intended deletion by PCR, with primers flanking and internal to the microdeletion (**Supplementary Table S2**).

### Taqman real-time PCR

Reverse transcription was performed using oligo d(T) and Superscript III first-strand synthesis system (Invitrogen) according to the manufacturer’s conditions. TaqMan real-time PCR was carried out using the following probes for human genes: *FAAH* (Hs01038660_m1), *FAAH- OUT* (Hs04275438_g1), *BDNF* (Hs03805848_m1), *ACKR3* (Hs00604567_m1), *WNT5B* (Hs01086864_m1), *GABBR2* (Hs01554996_m1), *DKK1* (Hs00183740_m1), *SFRP2* (Hs00293258_m1), *SERPINF1* (Hs01106937_m1) and *ACTB* (Hs01060665_g1). Mouse Taqman probes used were: *Faah* (Mm01191801_m1) and *Actb* (Mm01205647_g1). The expression level of target genes was normalized to the housekeeping Actin gene mRNA. Relative gene expression (relative quantities [RQ] value) was determined using the 2^−ΔΔCt^ equation in which control unaffected individuals or empty vector cDNA samples were designated as the calibrator. All RT–PCR data are expressed as mean ±SEM with significance indicated by *p≤0.05, **p≤0.01 and ***p≤0.001 (two tailed Student’s t-test).

### siRNA assay

In a six-well tissue culture plate, 10^5^ HEK293 cells per well were seeded in 2 ml antibiotic-free normal growth media supplemented with FBS. Transfections were carried out using 30 pmols of MISSION esiRNA targeting human *FAAH* (EHU098921; Sigma Aldrich) or a universal negative control (SIC001) with Lipofectamine RNAiMax transfection reagent (ThermoFisher), according to the manufacturer’s recommendations. Forty-eight hours after transfection, RNA was isolated and TaqMan real-time PCR was carried out.

### Chromatin immunoprecipitation (ChIP)

Chromatin Immunoprecipitation (ChIP) assays were performed according to the manufacturer’s protocol using the Chromatrap Enzymatic ChIP-seq kit. Immunoprecipitations were performed overnight at 4°C using antibodies against H3K27ac (Abcam 4729), H3K4me1 (Abcam 8895), H3K4me3 (Abcam 8580), H3K9me3 (Abcam 8898), H3K27me3 (Active Motif 39157), DNMT1 (Active Motif 39204) and DNMT3A (Active Motif 39206). Rabbit IgG were used as control for ChIP and primers within a gene desert on chromosome 16 were used as a negative control for qPCR.

All ChIP experiments, which were performed in triplicate, were done using two independent chromatin preparations. The immunoprecipitated DNA and the input DNA were analysed by real-time PCR using the ΔΔCt method and the primers listed in **Supplementary Table S3**.

### EpiTect Methyl II PCR Assay

EpiTect Methyl II PCR Primer Assay (Qiagen) was performed according to the manufacturer’s protocol. Briefly, 250 ng of genomic DNA was used to set up the four independent restriction enzyme digests: (1) methylation-sensitive, (2) methylation-dependent, (3) methylation-sensitive and methylation-dependent double digest or (4) mock digest. Q-PCR was performed as per the manufacturer’s protocol, using commercially available primers for human *FAAH* (CpG Island 100530) (EPHS100530-1A, Qiagen). Methylation-sensitive (EPHS115450-1A) and methylation-dependent (EPHS115451-1A) digest control assays were performed to test the cutting efficiency of the restriction enzymes. Samples were analysed as recommended by the manufacturer (Qiagen).

### Cellular fractionation

Cytoplasmic and nuclear RNA were isolated with the Cytoplasmic & Nuclear RNA Purification Kit (Norgen Biotek; Cat. 21000) following the manufacturer’s manual. The effectiveness of cellular separation was controlled with cytoplasmic and nuclear markers *ACTB* and *U6*, respectively (see **Supplementary Table S4**).

### Fibroblast cell lines

Ethical approval was granted by University College London REC and written informed consent was provided by patient PFS and four gender-matched healthy controls. A punch skin biopsy (3-6mm) was taken from the outer upper arm of each individual and primary cultures of dermal fibroblasts were passaged in Dulbecco’s Modified Eagle’s Medium (Thermo Fisher) supplemented with 10% fetal bovine serum (Hyclone) and 1% penicillin/streptomycin (Thermo Fisher).

### Wound healing (scratch) assay

Primary fibroblast cultures derived from patient PFS and a healthy female control were seeded on 6-well plates and allowed to grow until confluent. The control fibroblast cell line, like PFS, was heterozygous for the hypomorphic SNP rs324420 but did not carry the ∼8 kb *FAAH-OUT* microdeletion. Fibroblasts were serum-starved for 24 h, and the bottom of the dish was scraped with a pipette tip (0.2 mL) to create a standardized cell-free area. Cell migration through the space was tracked with time-lapse microscopy in a Nikon BioStation CT (Nikon, Nikon Instruments Europe BV, Netherlands). There, the cells were cultured at 37°C and 5% CO_2_, and phase contrast images of wounded areas were automatically recorded every 12 h during the gap closure. The images were analysed by Fiji/ImageJ to computationally measure and plot the gap area as a function of time and then the time it took for the gap to close to half of its original area was calculated.

### Microarrays

Total RNA was isolated from the primary fibroblast cultures derived from patient PFS and 4 healthy unrelated gender-matched controls (3 homozygous for wild-type C allele at rs324420 and 1 heterozygous C/A) using the PureLink RNA Micro Kit (Invitrogen) and run by Eurofins on the human Clariom D transcriptomic array (Thermo Fisher) using the GeneChip WT Plus labelling kit reagent. Expression data was RMA normalized and analysed using the Transcriptome Analysis Console (TAC) software (Thermo Fisher) and Ingenuity Pathway Analysis (Qiagen). Microarray data has been deposited at Gene Expression Omnibus Array Express with reference number E-MTAB-11809.

### ELISA

All experiments were performed in accordance with the UK Animals (Scientific Procedures) Act 1986 with prior approval under a Home Office project licence (PPL 70/7382). Mice were kept on a 12-hr light/dark cycle and provided with food and water ad libitum.

The FAAH inhibitor URB597 (0.3 mg/kg; i.p.) (Cayman Chemical, MI, USA) was freshly dissolved in vehicle containing 5% dimethylsulfoxide (DMSO), 5% Tween-80 and 90% saline. Controls were given the vehicle only. Adult male C57BL/6 mice (Charles River) were intraperitoneally injected with either URB597 (0.3 mg/kg, n = 3) or vehicle (vol:1 ml/kg, n = 3). Twenty-four hours later, mice were euthanized by CO_2_ asphyxiation followed by cervical dislocation. The hippocampus was dissected and BDNF content measured using a commercially available sandwich enzyme-linked immunosorbent assay (ELISA) kit (Quantikine ®ELISA-Total BDNF, R&D Systems, Minneapolis, MN, USA) according to the manufacturer’s instructions.

### RNAscope *In situ* Hybridization (ISH)

Human Dorsal Root Ganglion (DRG) Paraffin Sections (HP-240, 5 µm thick), Human Brain Cerebral Cortex Frozen Sections (HF-210, 7-10 µm thick), Human Cerebellum Frozen Sections (HF-202, 7-10 µm thick) and Human Prostate Frozen Sections (HF-408, 7-10 µm thick) were obtained commercially from Zyagen (www.zyagen.com) via AMS Biotechnology (https://www.amsbio.com) (**Figure 2** and **Supplementary Figures S1 - S2**).

For the mouse DRG sections (**Supplementary Figure S3**), adult C57BL/6 mice were deeply anesthetized with pentobarbital (i.p.) and transcardially perfused with heparinized saline (0.9% NaCl) followed by 25 ml of cold 4% paraformaldehyde in phosphate-buffered saline (pH 7.4). DRGs were extracted from the lumbar area and post-fixed with the same fixative solution for 2 hours at 4 °C before being embedded in cryopreservative solution (30% sucrose) overnight at 4°C. Tissue samples were then placed in OCT blocks for posterior sectioning by cryostat. 11 μm thick sections were mounted onto Superfrost Plus (Fisher Scientific) slides, allowed to freeze-dry overnight at -80 °C, for an immediate use, or were stored at −80 °C in air-tight containers for no longer than a month for subsequent experiments.

*In situ* hybridization was performed using the RNAscope assay (Advanced Cell Diagnostics) following the protocol for fresh-frozen samples, in the case of Human Cerebral Cortex, Human Cerebellum and Human Prostate tissue samples, and mouse DRG samples with 1h post-fixing with 4% PFA in PBS at 4 °C and stepwise dehydration with 50%, 70% and 100% Ethanol. Tissue pre-treatment consisted of hydrogen peroxide and Protease IV (10 and 20 min respectively) at RT. Following pre-treatment, probe hybridization and detection with the Multiplex Fluorescence Kit v2 were performed according to the manufacturer’s protocol.

Human DRG Paraffin Sections were treated according to the ACD’s FFPE-fixed samples protocol.

Probes included *hsNEFH* (#448141-c4), *hsCNR1* (#591521-c4)*, hsFAAH* (#534291-c2) and *hsFAAH-OUT* (#534301-c3). RNA localisation was detected with either AF488 or Opal 520 (green), Opal570 (red) and Opal 650 (far-red) fluorochrome dyes (Perkin Elmer) compared to DAPI staining (nuclei) or TS-coumarin (TS405, Perkin Elmer) used for *NEFH or CNR1*. ISH slides were mounted using Prolong Gold (ThermoFisher Scientific #P36930). Mouse RNAscope probes included *mmFaah* (#453391) *and mmNefh* (#443671-c4).

Fluorescence was detected using Zeiss LSM 880 Airyscan microscope. Images were taken at 10x and 20x magnifications with 4x averaging. Tiles were stitched when more than one was used to image the area, airyscan processed and exported as 16-bit uncompressed tiff files for further basic editing in Adobe Lightroom v6 (Adobe) on a colour calibrated iMac (X-Rite) retina monitor. Final images were exported as jpeg files with 7,200 pix on longest side at 300 ppi.

### Statistical analysis

Data was analysed using GraphPad Prism 9 (GraphPad Software, Inc), and results presented as mean ± SEM with n referring to the number of samples tested per group, as indicated in figure legends.

## Acknowledgements

We would like to thank patient PFS, Dr Devjit Srivastava and the volunteers who participated in this study. Addgene plasmid 61591 was a gift from Feng Zhang, Addgene plasmid 106219 was a gift from Charles Gersbach and Addgene plasmid 68495 was a gift from George Church.

## Author contributions

Conceptualization: HM, AMH, ALO, JJC

Methodology: HM, AMH, APL, AAF, HH, ALO, JJC

Investigation: HM, AMH, CY, SSV, APL, KP, SZ, AAF, HH, JNW, ALO, JJC

Visualization: HM, ALO,

Supervision: HH, JNW, ALO, JJC

Writing-original draft: HM, ALO, JJC

Writing-review&editing: HM, AMH, CY, SSV, APL, AAF, HH, JNW, ALO, JJC

## Data and materials availability

Microarray data has been deposited at Gene Expression Omnibus Array Express with reference number E-MTAB-11809. All data are available in the main text or supplementary materials.

**Supplementary Figure S1.**
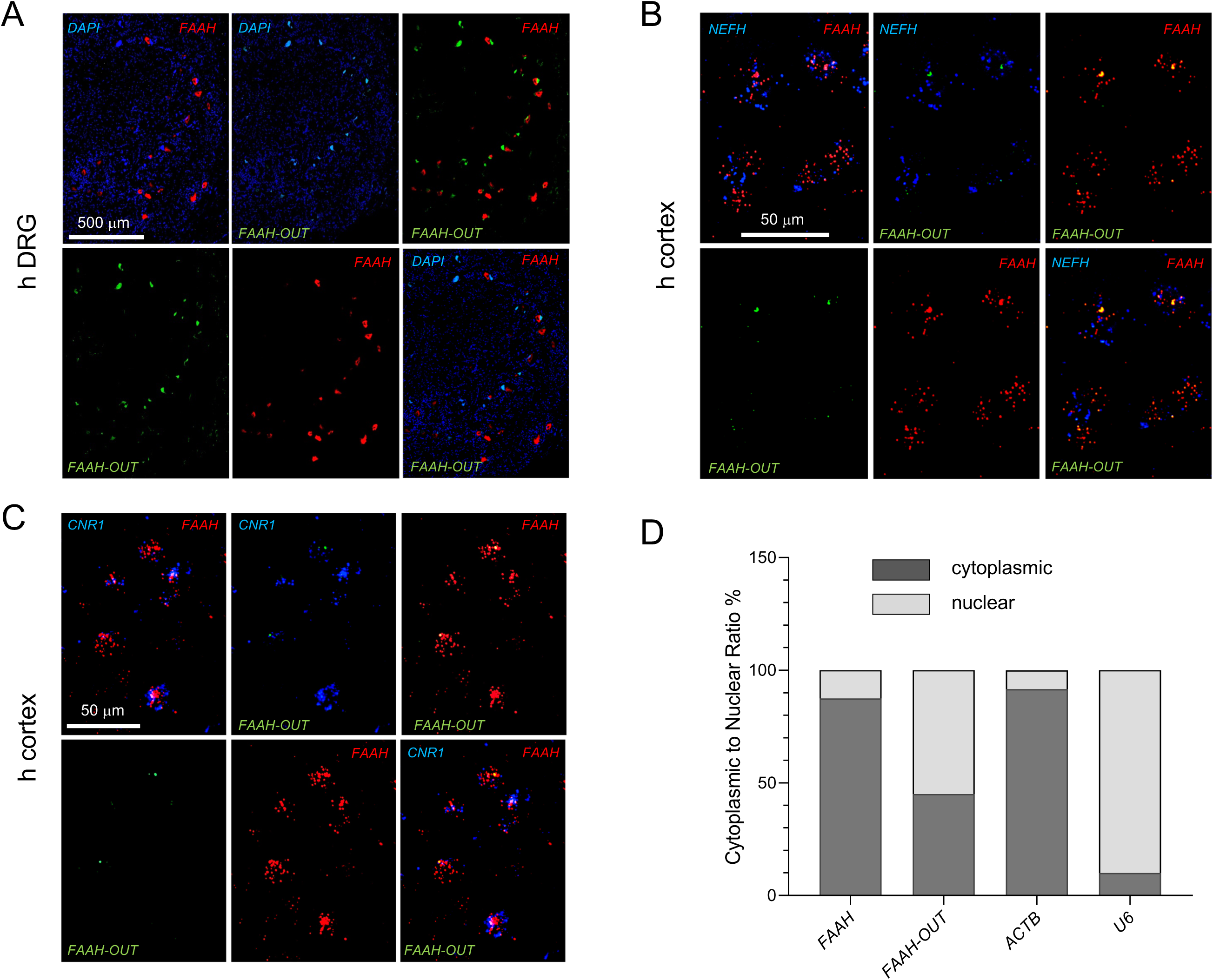
*FAAH* and *FAAH-OUT* RNA expression levels and localisation. A. *FAAH* and *FAAH-OUT* RNA expression levels and localisation in human DRG neurons. 5 μM -thick cuts of human DRGs were analysed by RNAscope assay. Localisation of *FAAH-OUT* lncRNA (green, AF488) compared to *FAAH* mRNA (red, Opal570) as analysed by RNAscope assay in 5 μM - thick cuts of human DRGs. DAPI staining indicates nuclei (blue). Scale bars are shown in white. **B** and **C. *FAAH* and *FAAH-OUT* RNA expression levels and localisation in human cerebral cortex neurons**. 7-10 μM - thick fresh-frozen cuts of human cerebral cortex were fixed with 4% PFA and analysed by RNAscope assay. **B** – four representative cells show localisation of *FAAH-OUT* lncRNA (green, AF488) compared to *FAAH* mRNA (red, Opal570) and *NEFH* mRNA (blue, TS405) localisation. **C** – five representative cells show localisation of *FAAH-OUT* lncRNA (green, AF488) compared to *FAAH* mRNA (red, Opal570) and *CNR1* mRNA (blue, TS405) localisation. Scale bars are shown in white. **D. Subcellular localisation of *FAAH* and *FAAH-OUT* RNAs.** RT-qPCR analysis of RNA following nuclear/cytoplasmic fractionation shows the distribution of the indicated transcripts in HEK293 cells. *FAAH* mRNA is predominantly cytoplasmic whereas *FAAH-OUT* lncRNA is enriched in the nucleus. *ACTB* and *U6* RNAs were used as cytoplasmic and nuclear controls respectively. The RT-qPCR data, represented as a percentage of the total amount of detected transcripts, are presented as mean of technical triplicates.

**Supplementary Figure S2A.**
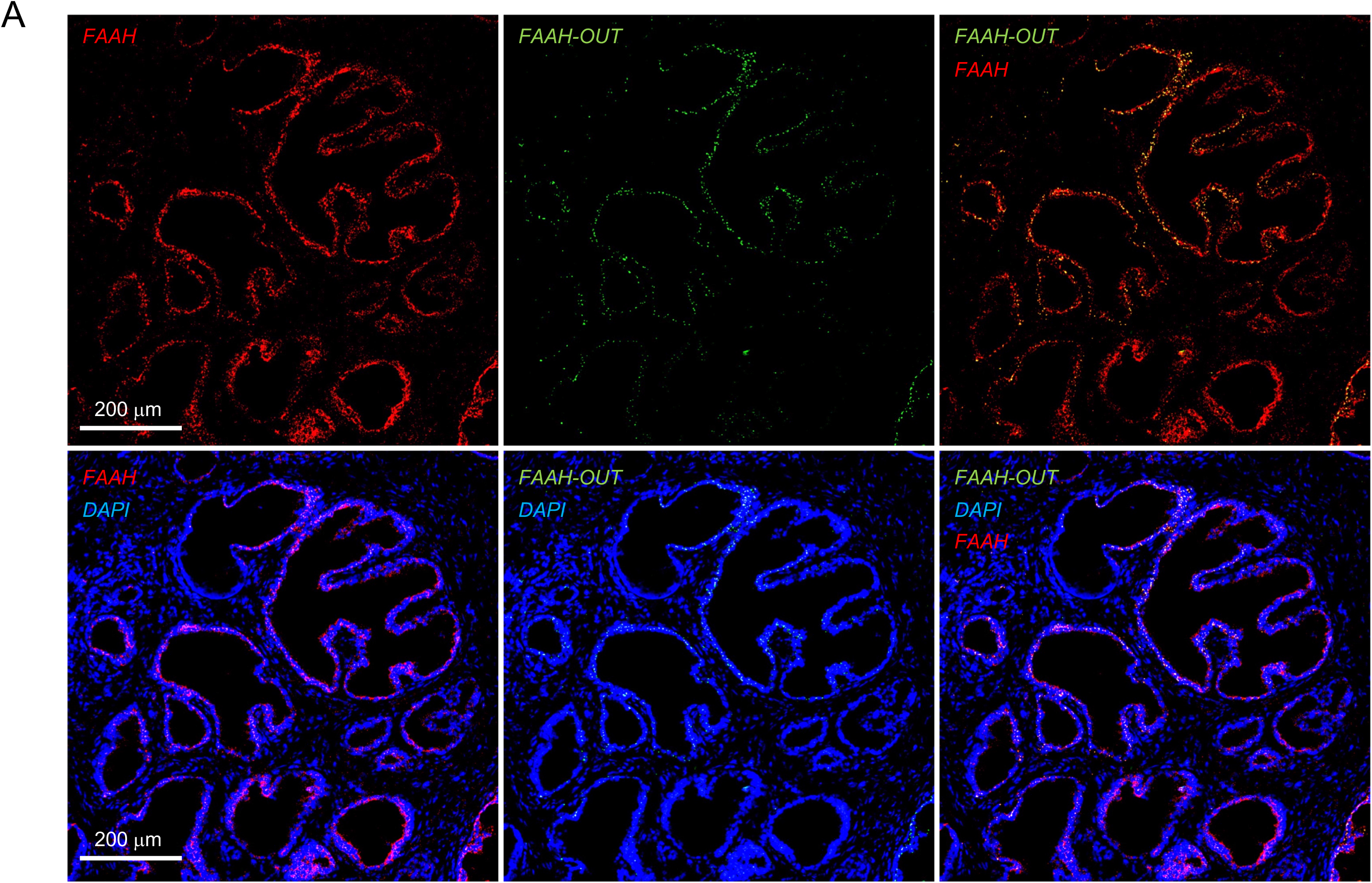
*FAAH* and *FAAH-OUT* RNA expression levels and localisation in human prostate tissue cells. 7-10 μM - thick cuts of fresh frozen prostate sections were analysed by RNAscope assay (Methods). Localisation of *FAAH-OUT* lncRNA (green, AF488) was compared to *FAAH* mRNA (red, Opal570) localisation and DAPI staining indicating nuclei positions (blue). Scale bars (200 mM) are in white.

**Supplementary Figure 2B and C.**
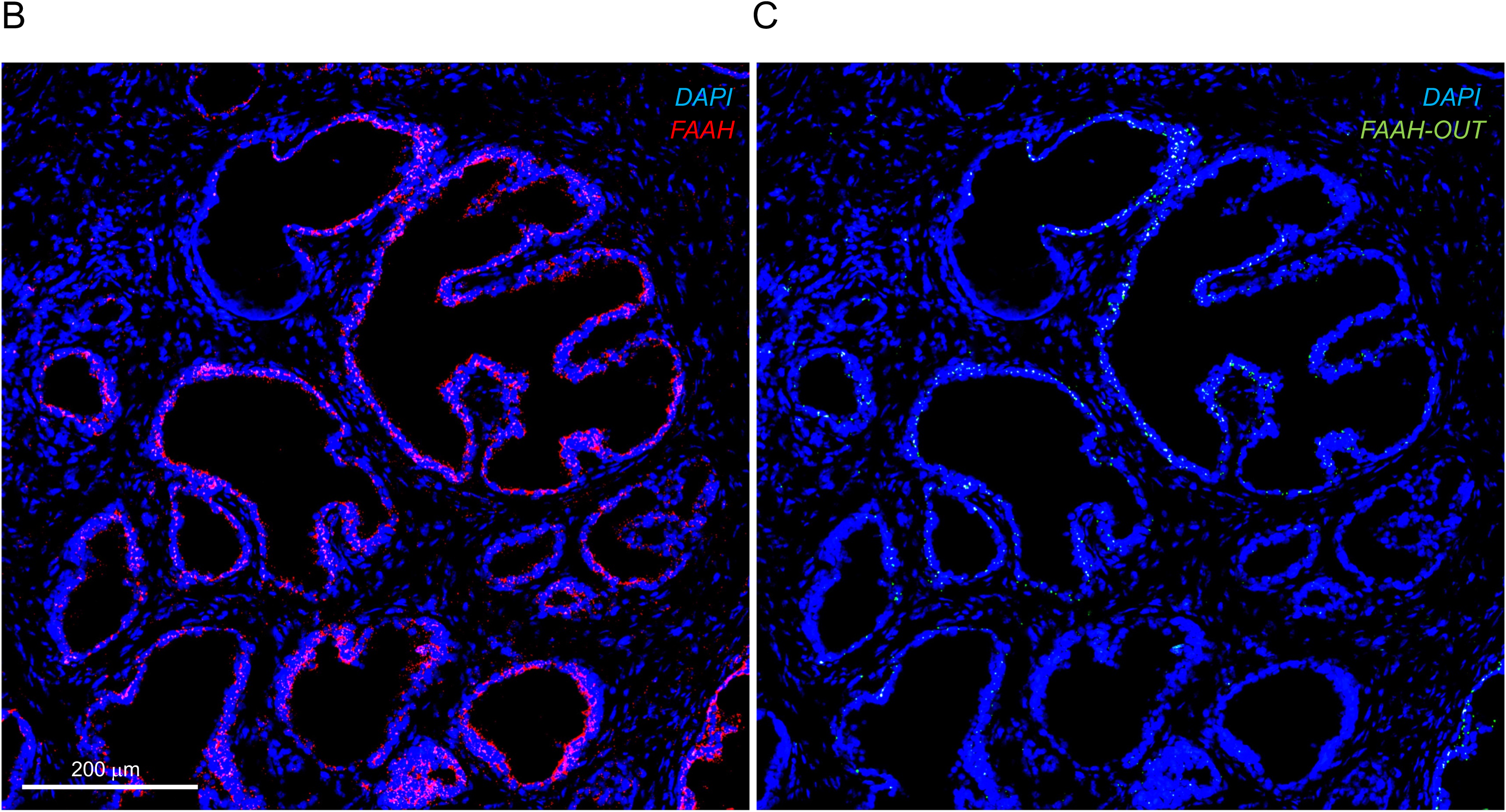
*FAAH* and *FAAH-OUT* RNA expression levels and localisation in human prostate tissue cells. **B** and **C**. Enlarged views of prostate tissue section showing *FAAH* vs DAPI and *FAAH-OUT* vs DAPI and demonstrating mostly nuclear localisation for *FAAH-OUT* lncRNA and mostly cytoplasmic localisation for *FAAH* mRNA. Scale bar (200 mM) is in white.

**Supplementary Figure S3.**
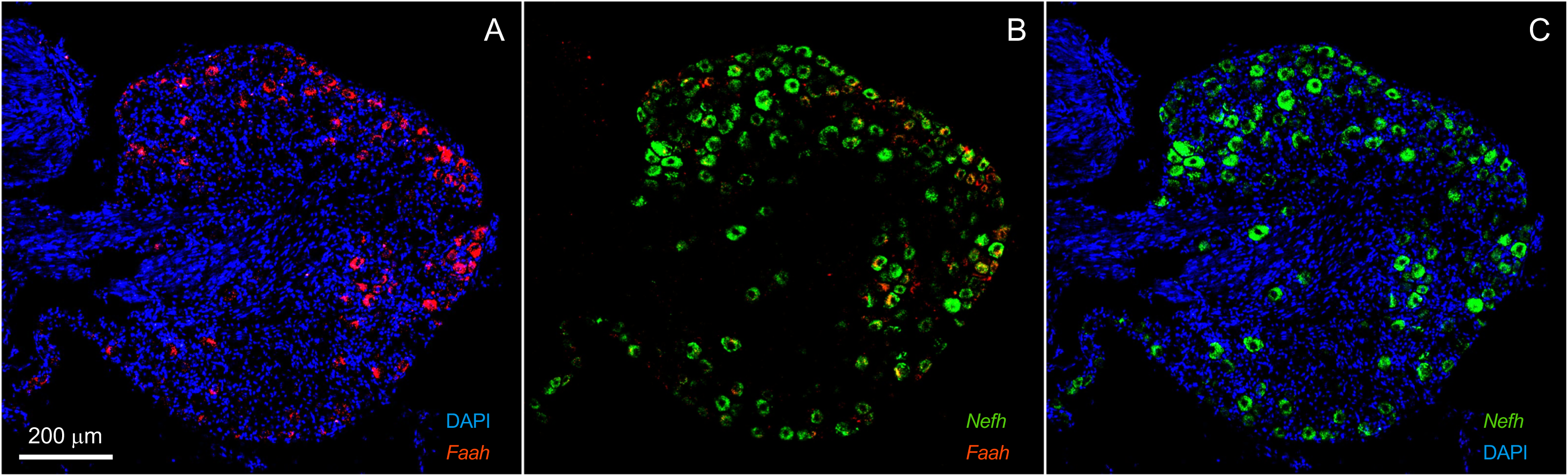
*Faah* RNA expression levels and localisation in mouse DRG *Faah* RNA expression levels and localisation in mouse DRG neurons. 11 μM - thick fresh-frozen cuts of mouse lumbar DRGs were fixed with 4% PFA and analysed by RNAscope assay. **A**-**C**. Localisation of *Faah m*RNA (**A**, red, Opal570) compared to *Nefh* mRNA (**B** and **C,** green, AF488) localisation and DAPI staining indicating nuclei positions (**A** and **C**, blue). Scale bar (200 mM) is shown in white.

**Supplementary Figure S4A.**
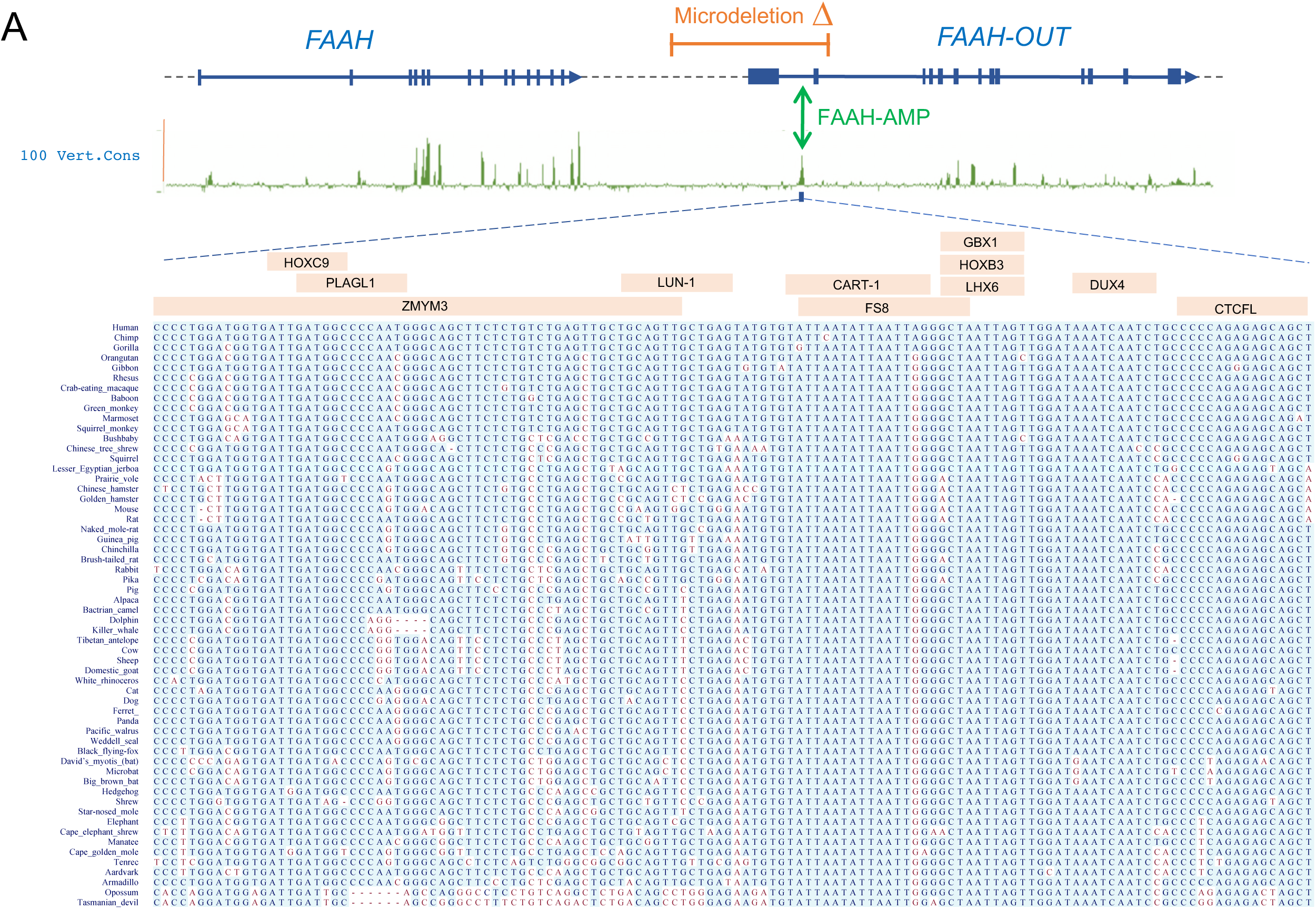
FAAH-AMP is highly conserved among species. Map showing relative positions of the ∼8 kb microdeletion identified in patient PFS (in orange) and ‘FAAH-AMP’ conserved element indicated with green double arrow. Exons are denoted by blue boxes and the direction of transcription shown by arrows. The PhyloP base-wise conservation track for 100 vertebrates from the UCSC genome browser shows regions of high sequence conservation as peaks (in green), with the majority of these mapping to gene exons in *FAAH* and *FAAH-OUT*. The core of the conserved element (chr1:46,424,925-46,425,054; build hg38) is expanded in multiple alignment of the representative vertebrates. The positions of DNA-binding for selected human transcription factors (TFs) are indicated according to Transfac and Jaspar databases.

**Supplementary Figure S4B.**
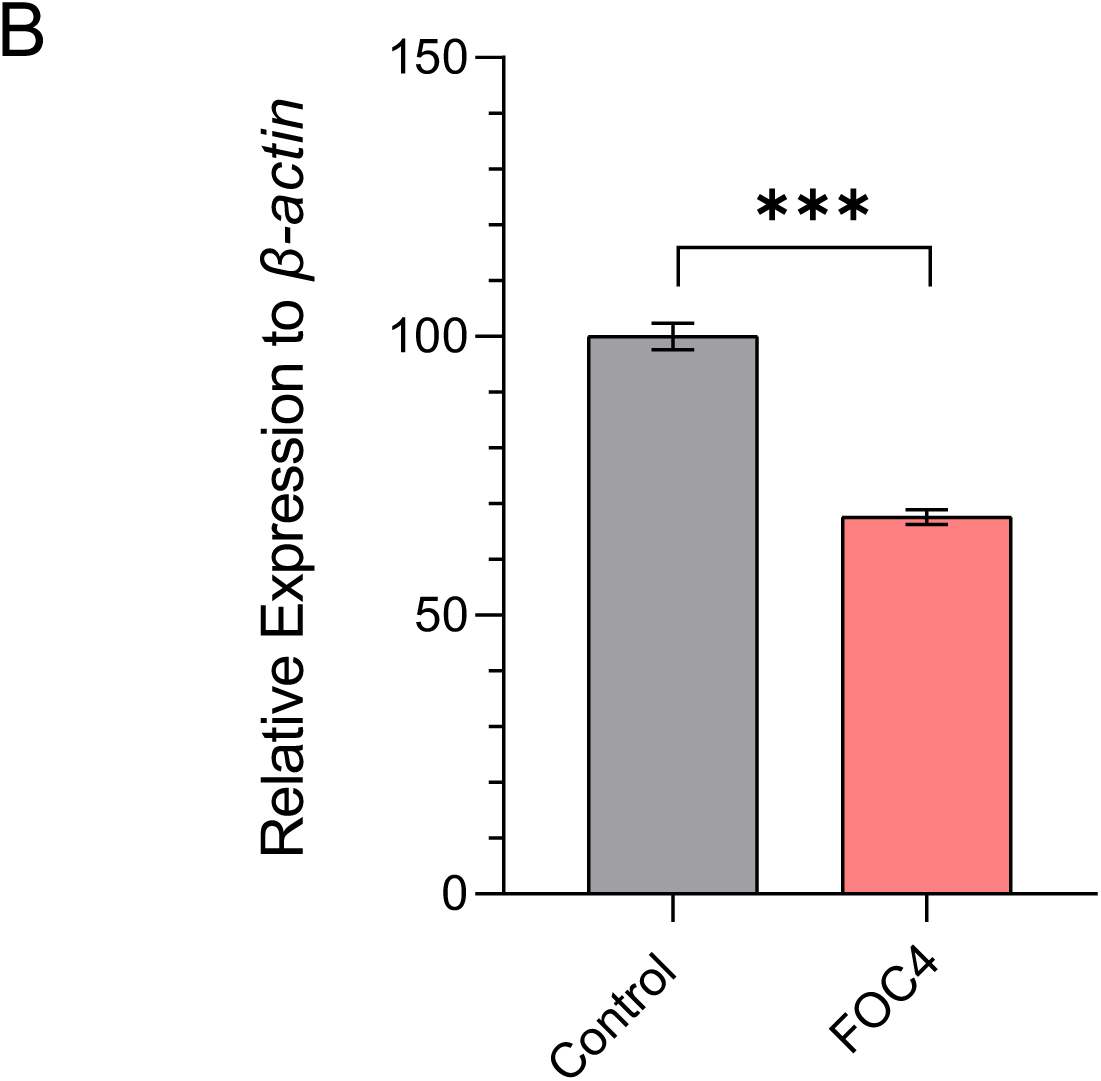
CRISPR/Cas9-induced deletion of Faah-AMP leads to reduction in *Faah* expression in mouse cells. RT-qPCR analysis of *Faah* mRNA levels showed a significant reduction when mouse CAD cells were transiently transfected with a CRISPR/Cas9 construct containing the FOC4 guide RNA pair, which is designed to delete the Faah-AMP conserved element. The normalized expression value of control (empty vector) was set to 100 and all other gene expression data were compared to that sample. Data are expressed as mean of triplicates ± SEM and analysed by Student’s t-test, *** *P* ≤0.001.

**Supplementary Figure S5.**
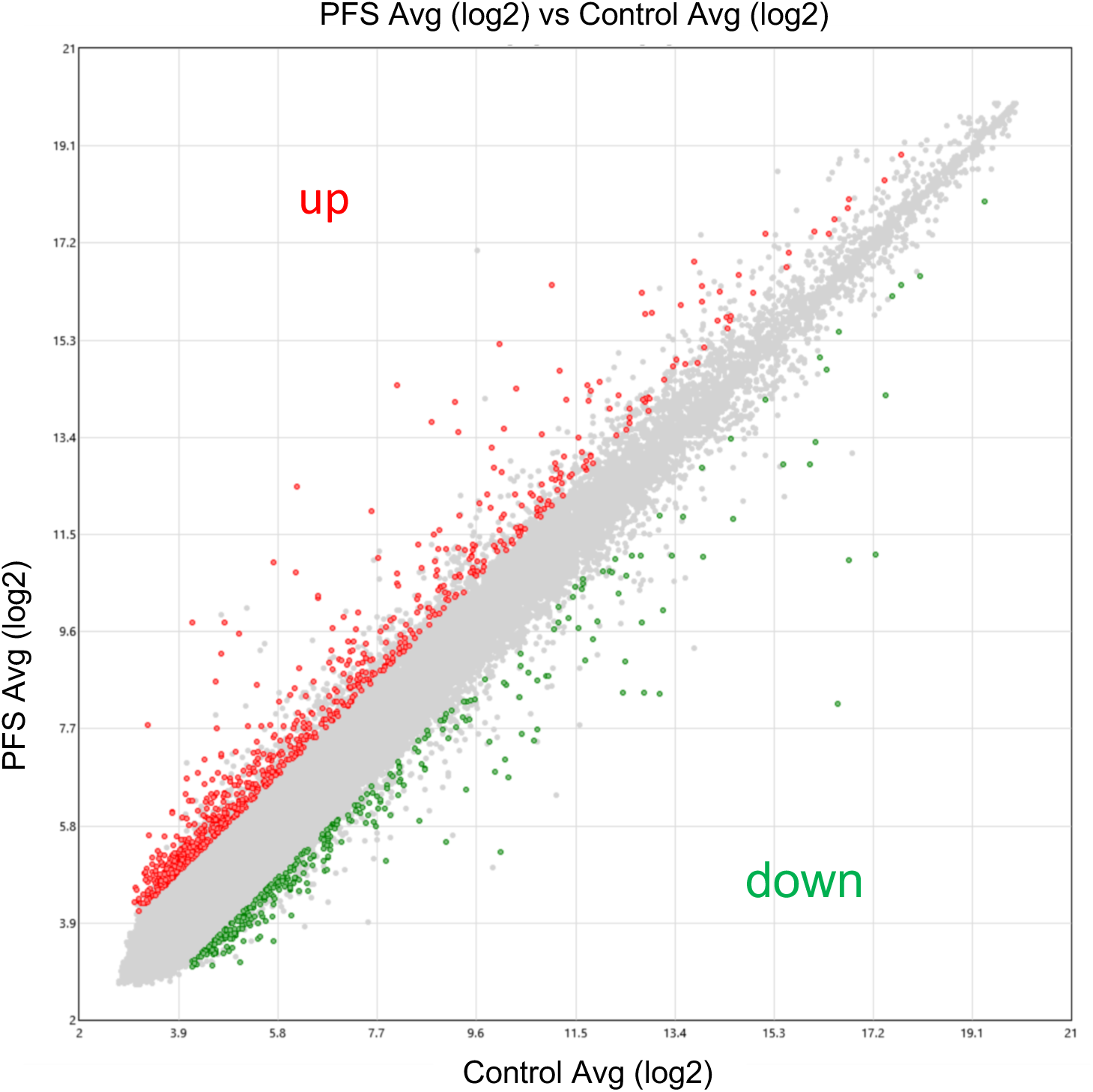
Scatter plot of microarray data showing gene expression changes in PFS-derived fibroblasts vs 4 controls. Scatter plot of normalized probe intensities (log2) for significantly dysregulated genes from the Clariom D microarray analysis of PFS-derived fibroblasts (y axis) vs 4 gender-matched fibroblast controls (x-axis). Each circle denotes a single gene with red circles showing up-regulated genes in PFS (>2 fold; p<0.05) and green circles showing down-regulated genes in PFS (>2 fold; p<0.05)

**Supplementary Figure S6.**
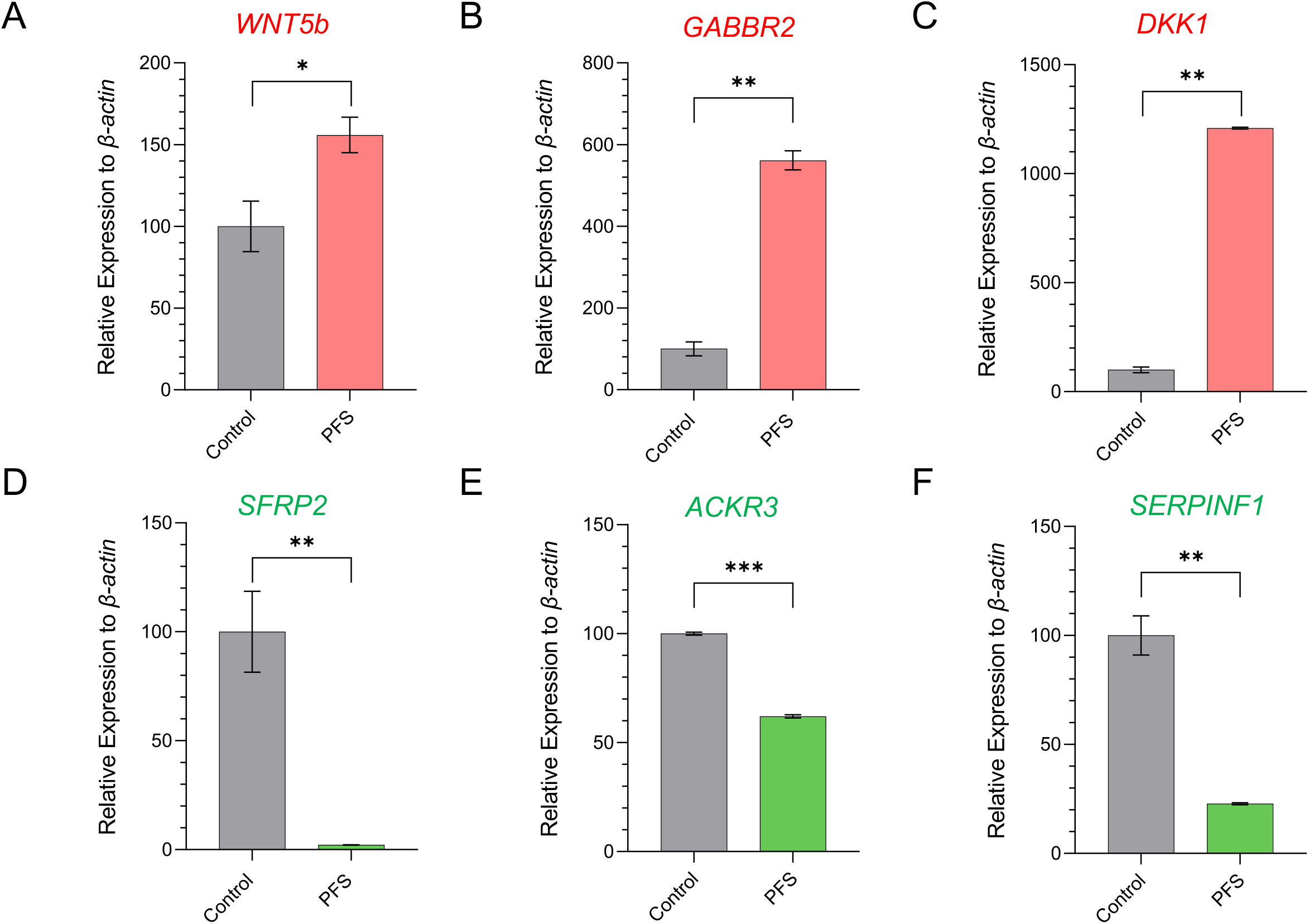
Differential expression of key genes in patient fibroblasts confirmed by RT-qPCR. RT-qPCR analysis of key Differentially Expressed Genes (DEGs) mRNA levels in patient (PFS) fibroblasts showed a significant rise for (**A**) *WNT5b* (1.5-fold); (**B**) *GABBR2* (5.6-fold) and (**C**) *DKK1* (12-fold) expression when compared to gender matched controls. The data also showed significant reduction in expression levels of (**D**) *SFRP2* (47.5-fold); (**E**) *ACKR3* (1.6-fold) and **(F)** *SERPINF1* (4.4-fold) when compared to the controls. Data were normalized to the beta-actin gene as an endogenous control. The normalized expression value of control subjects was set to 100. Data are expressed as mean of triplicates ± SEM and analysed by Student’s t-test, \**P* ≤ 0.05, \*\**P* ≤ 0.01, *** *P* ≤ 0.001

**Supplementary Figure S7A.**
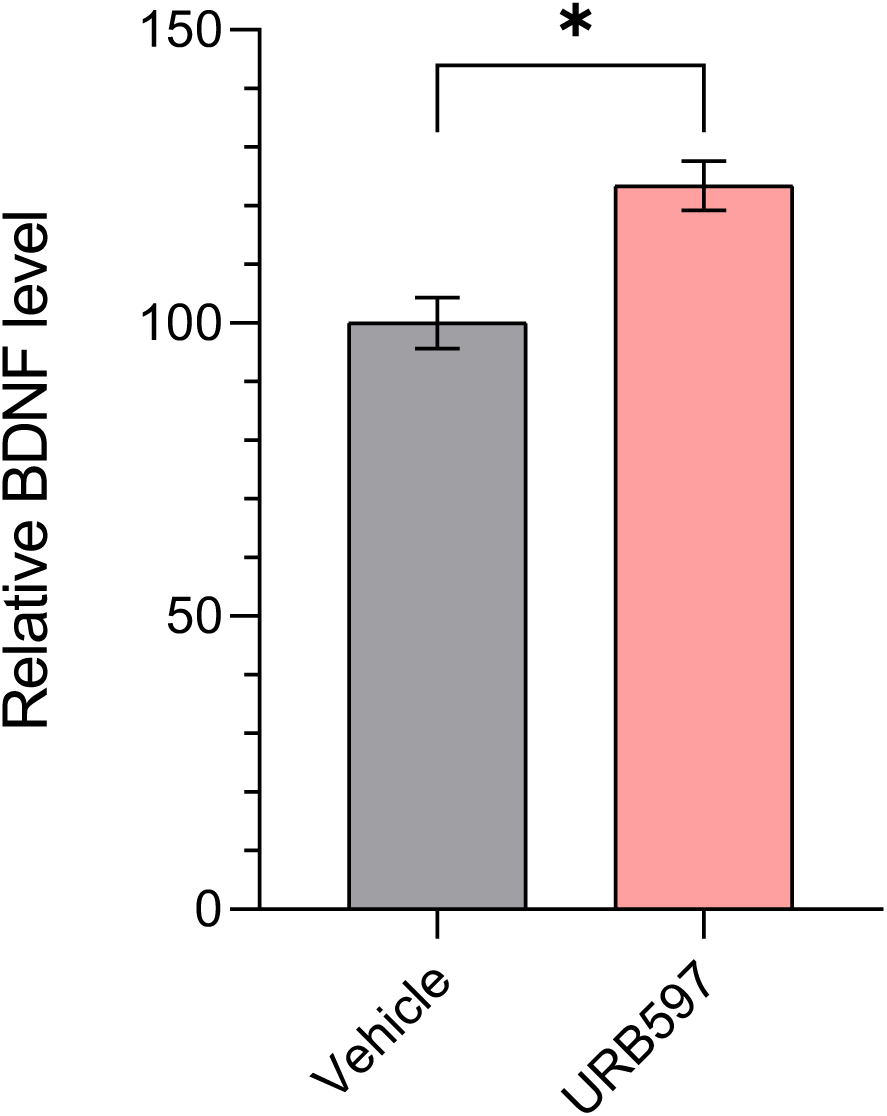
Effects of pharmacological inhibition of *FAAH*. Mice treated with FAAH inhibitor URB597 showed significantly higher BDNF (brain-derived neurotrophic factor) levels in the hippocampus (detected by ELISA) compared with vehicle treated mice (*n* = 3 per group; Student’s t-test, \**P* ≤ 0.05).

**Supplementary Figure 7B.**
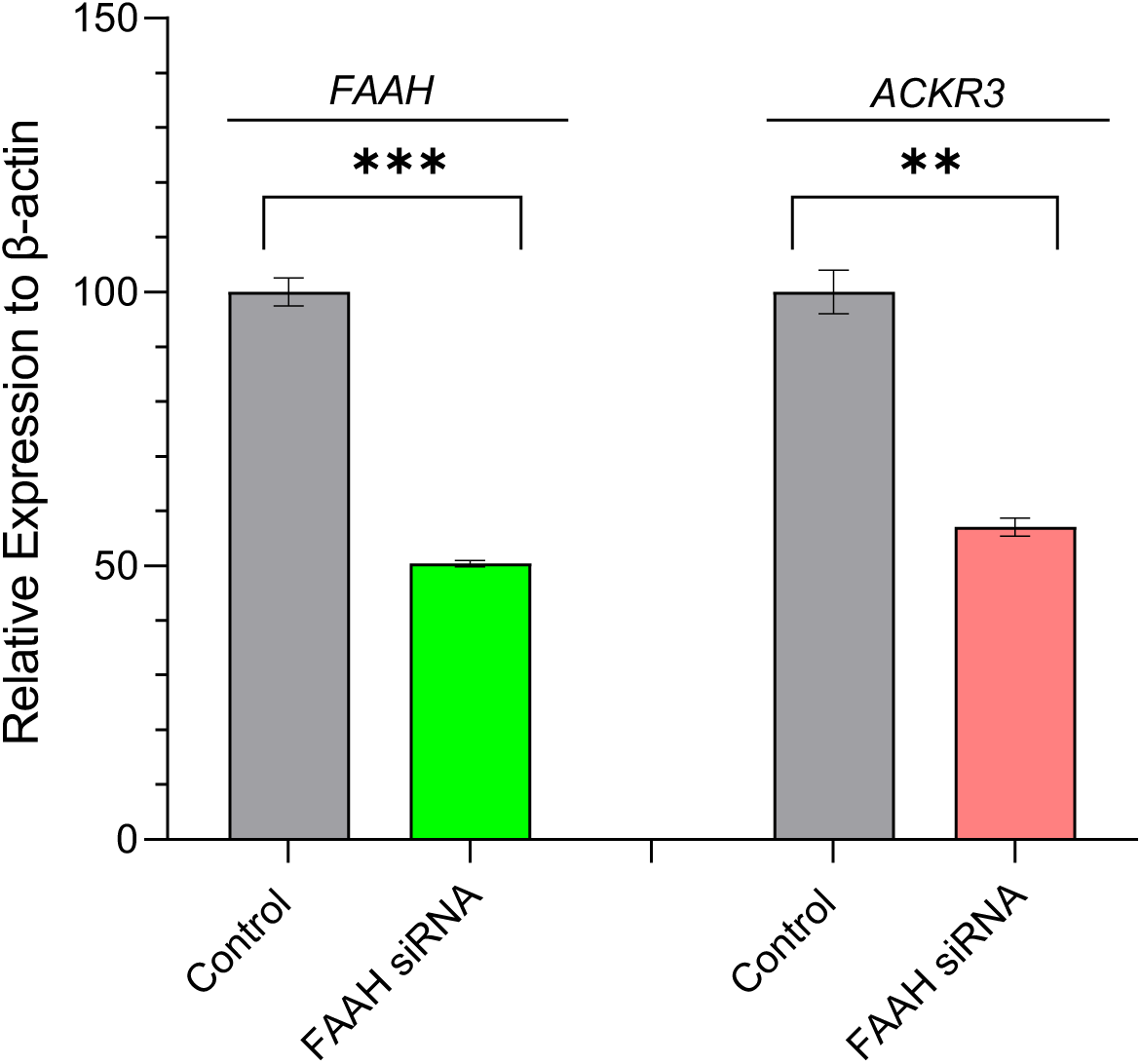
Effects of siRNA knockdown of *FAAH*. *FAAH* and *ACKR3* expression in HEK293 cells following *FAAH* siRNA treatment. Expression levels of both genes are significantly reduced after *FAAH* knockdown. Data are expressed as mean of triplicates ± SEM and analysed by Student’s t-test; \*\**P* ≤ 0.01, *** *P* ≤ 0.001.

**Supplementary Figure S8.**
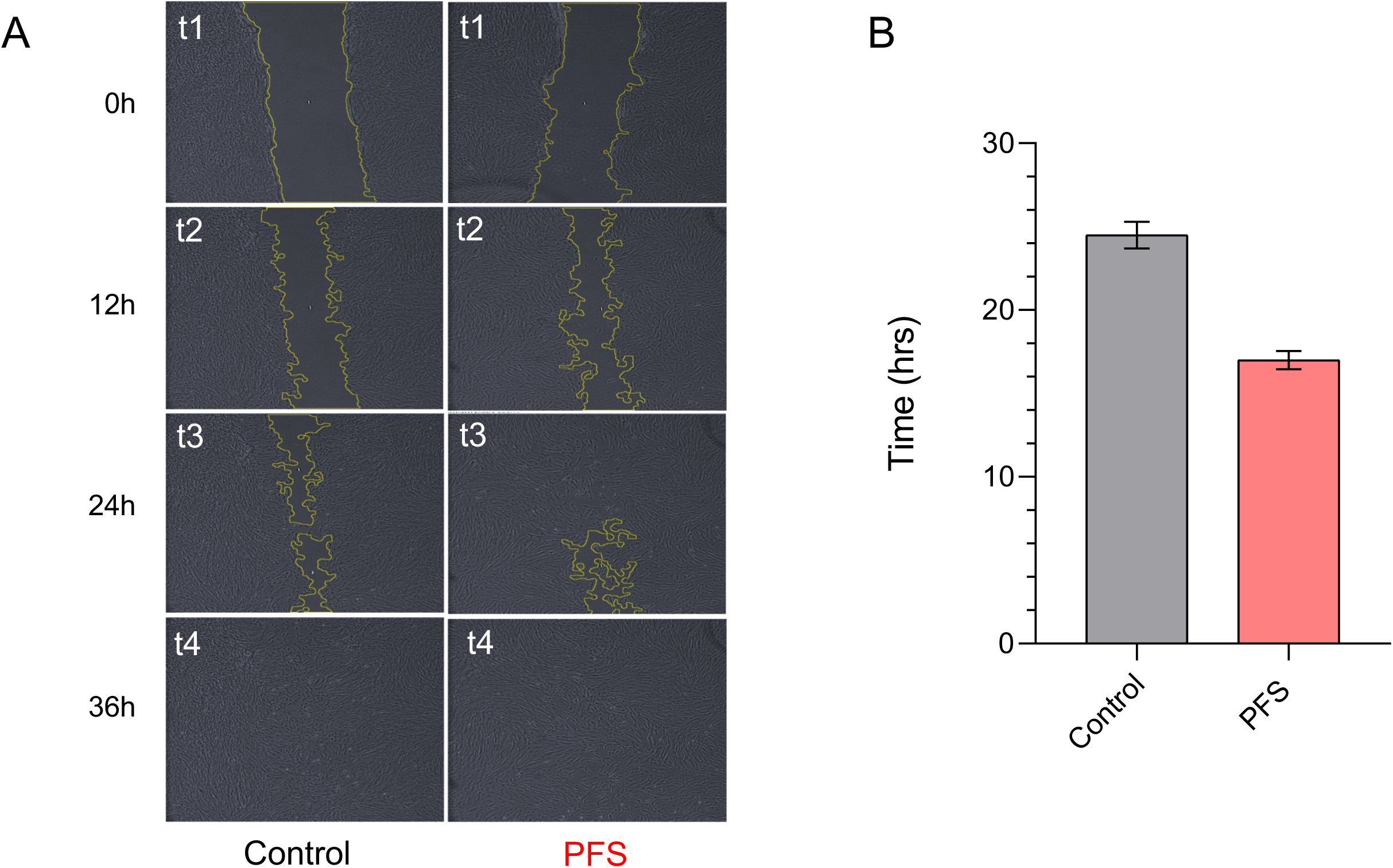
The microdeletion in *FAAH-OUT* leads to an acceleration of wound healing. **A**. Wound healing scratch assay was performed using fibroblasts from the PFS patient and a gender matched control who carries the hypomorphic SNP (rs324420) but not the *FAAH-OUT* microdeletion. Images of the recovering monolayer of cells were taken from the start of the experiment (t1=0h) and the following intervals of 12 h for the next 72h (t1, t2, t3, t4 time points correspond to 0, 12, 24 and 36h respectively). **B.** The images were analysed by Fiji/ImageJ to measure and plot the recovery of the initial gap area (resulting from the scratch) as a function of time. PFS fibroblasts recovered 50% of the scratched gap area in 17h time whereas control fibroblast needed 24.5h to achieve the same, resulting in approximately 30% faster recovery time for PFS fibroblasts.

**Supplementary Table S1:**
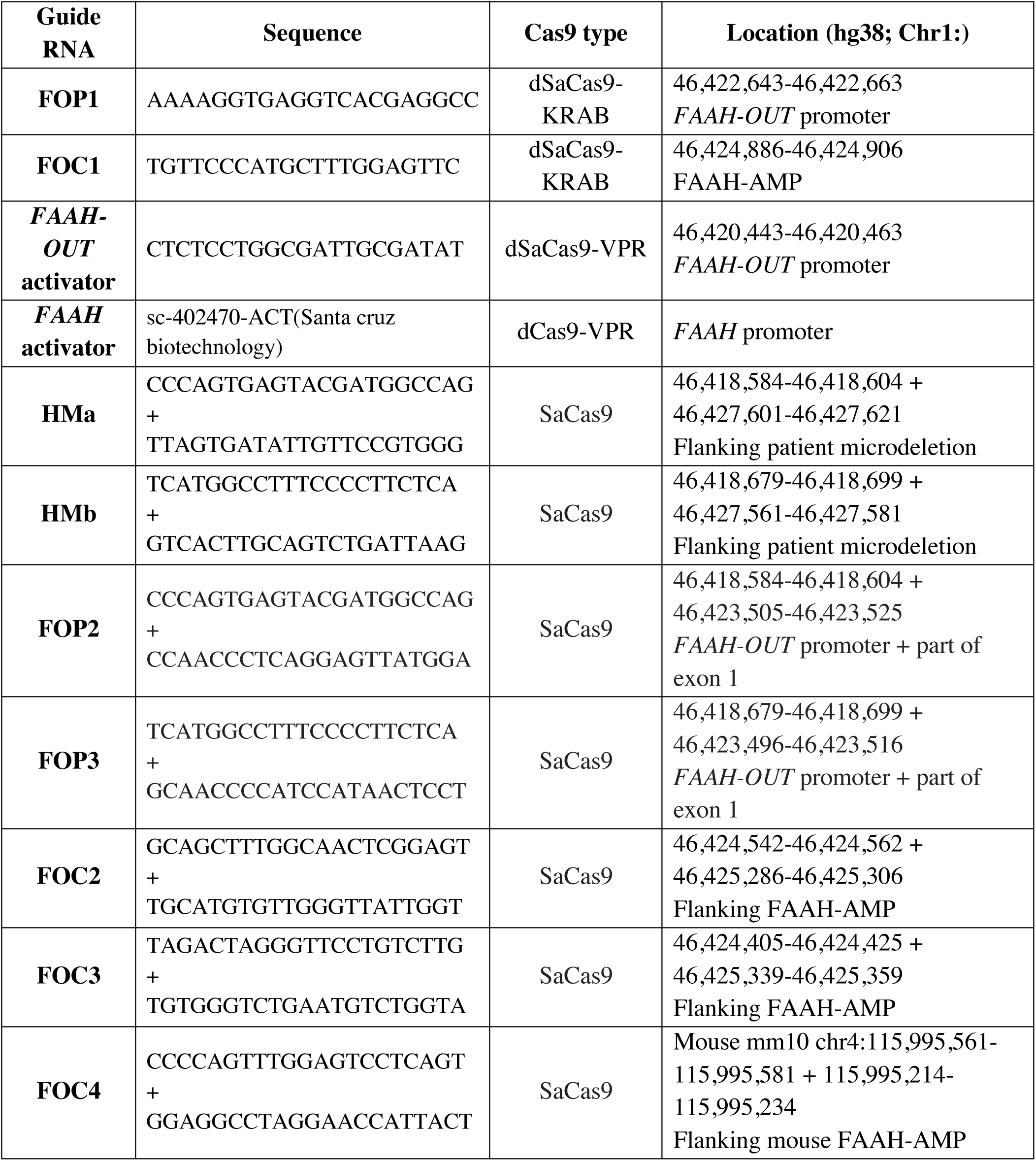
sgRNA sequences

**Supplementary Table S2:**
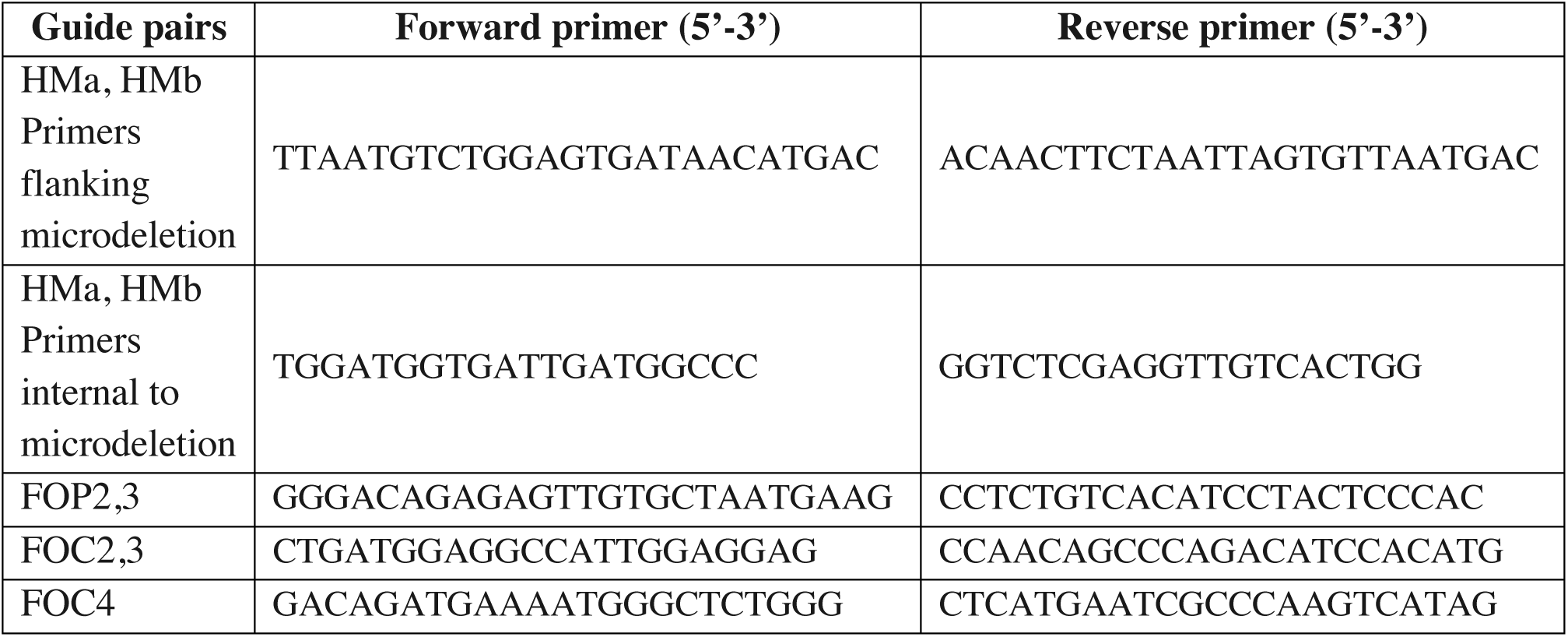
Primers used to confirm SaCas9-induced deletions

**Supplementary Table S3:**
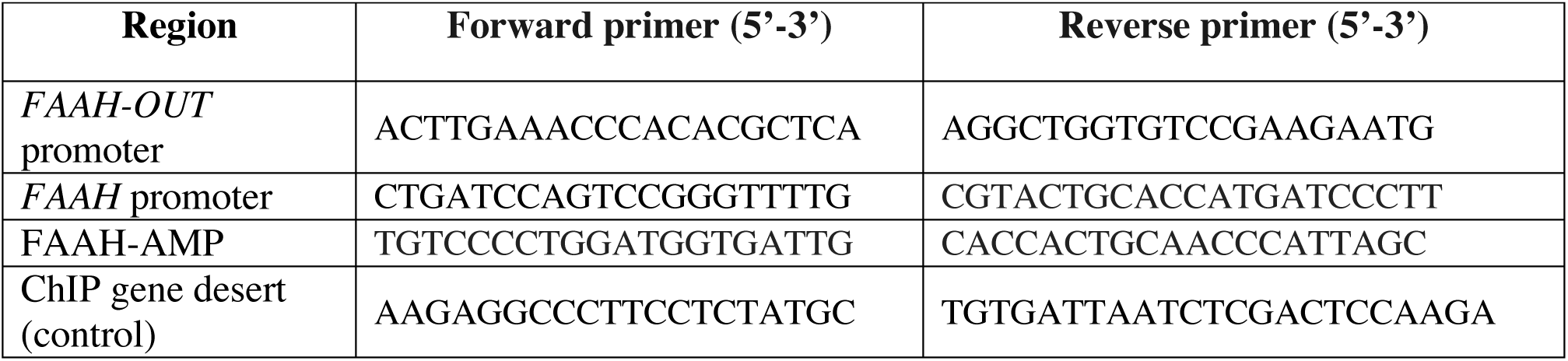
Primers used for ChIP-qPCR

**Supplementary Table S4:**
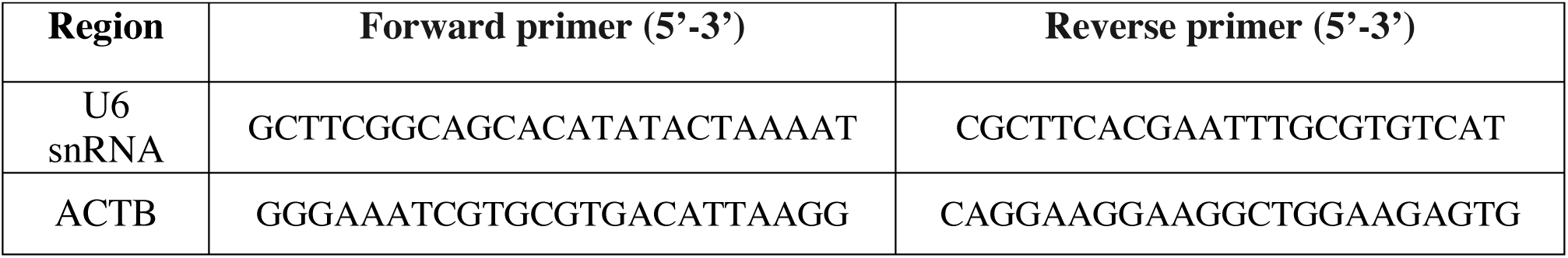
Primers used for cellular fractionation

